# Spike rate inference from mouse spinal cord calcium imaging data

**DOI:** 10.1101/2024.07.17.603957

**Authors:** Peter Rupprecht, Wei Fan, Steve J. Sullivan, Fritjof Helmchen, Andrei D. Sdrulla

**Affiliations:** Laboratory of Neural Circuit Dynamics, Brain Research Institute, University of Zurich, Switzerland; Neuroscience Center Zurich, University of Zurich, Switzerland; Department of Anesthesiology and Perioperative Medicine, Oregon Health & Science University, Portland, OR, USA; University Research Priority Program (URPP), Adaptive Brain Circuits in Development and Learning, University of Zurich, Switzerland

## Abstract

Calcium imaging is a key method to record the spiking activity of identified and genetically targeted neurons. However, the observed calcium signals are only an indirect readout of the underlying electrophysiological events (single spikes or bursts of spikes) and require dedicated algorithms to recover the spike rate. These algorithms for spike inference can be optimized using ground truth data from combined electrical and optical recordings, but it is not clear how such optimized algorithms perform on cell types and brain regions for which ground truth does not exist. Here, we use a state-of-the-art algorithm based on supervised deep learning (CASCADE) and a non-supervised algorithm based on non-negative deconvolution (OASIS) to test spike rate inference in spinal cord neurons. To enable these tests, we recorded specific ground truth from glutamatergic and GABAergic somatosensory neurons in the superficial dorsal horn of spinal cord in mice of both sexes. We find that CASCADE and OASIS algorithms that were designed for cortical excitatory neurons generalize well to both spinal cord cell types. However, CASCADE models re-trained on our ground truth further improved the performance, resulting in a more accurate inference of spiking activity from spinal cord neurons. We openly provide re-trained models that can be applied to spinal cord data of variable noise levels and frame rates. Together, our ground-truth recordings and analyses provide a solid foundation for the interpretation of calcium imaging data from spinal cord dorsal horn and showcase how spike rate inference can generalize between different regions of the nervous system.

**Significance Statement:** Calcium imaging is a powerful method for measuring the activity of genetically identified neurons. However, accurate interpretation of calcium transients depends on having a detailed understanding of how neuronal activity correlates with fluorescence. Such calibration recordings have been performed for cerebral cortex but not yet for most other CNS regions and neuron types. Here, we obtained ground truth data in spinal cord by conducting simultaneous calcium and electrophysiology recordings in excitatory and inhibitory neurons. We systematically investigated the transferability of cortical algorithms to spinal neuron subpopulations and generated inference algorithms optimized to excitatory and inhibitory neurons. Our study provides a foundation for the rigorous interpretation of calcium imaging data from spinal cord.

**Conflict of interest statement:** The authors declare no competing financial conflicts of interest.

## Introduction

The development of genetically encoded calcium indicators and two-photon microscopy have significantly improved our ability to measure neuronal population dynamics in intact tissues, providing unique insights into brain function over large spatiotemporal scales (Grienberger and Konnerth, 2012; Ji et al., 2016; Nelson et al., 2019). Calcium indicators, e.g., from the GCaMP family (Tian et al., 2009; Chen et al., 2013; Zhang et al., 2023), can be expressed using cell-specific promoters, allowing the characterization of neuronal activity in normal and pathological states in distinct neuronal populations. A fundamental problem remains to infer from the fluorescence signals the underlying neuronal activity, since calcium indicators provide only an indirect measure of neuronal spiking (Kerr et al., 2005; Yaksi and Friedrich, 2006; Lütcke et al., 2010). This is a non-trivial issue, as the relationship between calcium signals and electrophysiological spikes is non-linear and may depend on parameters such as the respective calcium indicator, expression levels, brain areas, cell types, and recording noise. It is therefore not clear whether an algorithm adapted for a specific dataset will generalize well to datasets recorded under different conditions. Optimizing spike inference thus requires not only the development and training of algorithms but also ground truth recordings to retrain and re-evaluate the performance of these algorithms. Such ground truth recordings are experimentally challenging since they require simultaneous electrophysiological recordings and calcium imaging from the same neurons. Most openly accessible ground truth datasets comprise only mouse neocortical neurons, and whether algorithms trained with these datasets can be applied to other neuronal populations, such as in the spinal cord, remains unknown.

The spinal cord is a complex structure supporting sensory, motor, and autonomic functions (Häring et al., 2018; Sathyamurthy et al., 2018). Multiple distinct populations exist in the dorsal horn of the spinal cord, broadly divided into excitatory (glutamatergic) and inhibitory (GABAergic) neurons, which can be targeted based on specific promoters (Fan and Sdrulla, 2020; Larsson, 2017; Todd, 2017). These populations receive and integrate somatosensory afferents and play critical roles in disease states such as pathological pain and itch (Christensen et al., 2016; Koch et al., 2018; Moehring et al., 2018; Peirs et al., 2020; Sullivan and Sdrulla, 2022). Importantly, they reside in the superficial laminae of the dorsal horn and therefore are accessible for two-photon imaging (Johannssen and Helmchen, 2013; Xu and Dong, 2019; Harding et al., 2020b).

Two-photon imaging of the spinal cord in the living animal is well established (Kerschensteiner et al., 2005; Misgeld et al., 2007; Drdla et al., 2009; Johannssen and Helmchen, 2010; Dibaj et al., 2010; Farrar et al., 2012; Ran et al., 2016), but the degraded visual accessibility due to heavily myelinated tissue together with motion artifacts pose major challenges (Davalos et al., 2008; Laffray et al., 2011; Nishida et al., 2014; Sekiguchi et al., 2016). Partly because of this reduced accessibility, only very few calcium recordings have been performed in conjunction with simultaneous electrophysiology (Harding et al., 2020a). A small number of studies have been performed to study the specific role of calcium signaling in spinal cord neurons (Harding et al., 2020a; Simonetti et al., 2013) but there are, to our best knowledge, no attempts to systematically calibrate calcium signals of spinal cord neurons for spike inference. This is problematic because spinal cord neurons differ from cortical neurons in their development and function, and it is therefore unclear how spike inference and calibration measurements from cortical calcium signals can be applied to spinal cord neurons.

Here, we conducted simultaneous cell-attached recordings and calcium imaging of dorsal horn neurons to study the relationship between calcium signals and electrophysiological action potentials. With this ground truth, we can systematically evaluate and re-train algorithms for spike rate inference, enabling a better-founded interpretation of calcium imaging data from spinal cord in the future.

## Materials and Methods

### Mouse Strains

VGlut2-Cre mice (Stock No: 016963, The Jackson Laboratory) were crossed with homozygous floxed GCaMP6s mice (derived from strain Ai96; Stock No: 024106, The Jackson Laboratory, Bar Harbor, ME), resulting in GCaMP6s expression in glutamatergic neurons. To drive expression of GCaMP6s in GABAergic interneurons, we crossed Viaat-Cre mice (Stock No: 016962, The Jackson Laboratory) with homozygous GCaMP6s mice. The mice were kept under standard colony conditions, with 12-hour day/night cycles, and had access to food and water ad libitum. Both males and females were used. All experiments were approved by the Institutional Animal Care and Use Committee at Oregon Health & Science University.

### *Ex vivo* lumbar spinal cord preparation

Four-to six-week-old Viaat/GCaMP6s or VGlut2/GCaMP6s mice of both sexes were deeply anesthetized with 5% isoflurane and decapitated. The lumbar spinal cord was rapidly removed *en bloc* and placed in oxygenated artificial cerebrospinal fluid (ACSF; in mM: 125 NaCl, 2.5 KCl, 26 NaHCO_3_, 1.25 NaH_2_PO_4_H_2_O, 1 MgCl_2_, 2 CaCl_2_, and 25 glucose) at room temperature. The dorsal roots were trimmed, and the dura was removed. During experiments, the tissue block was glued to a thick glass rectangle and perfused with room-temperature, oxygenated ACSF at 3 mL/min. In a subset of experiments (5 out of 21 recordings in VGlut2 mice), ACSF was warmed to physiological temperatures (37°C) to test for the effect of temperature on calcium indicator kinetics.

### Calcium imaging acquisition

For calcium imaging of spinal cord neurons *ex vivo* or *in vivo*, the imaging system consisted of a Zeiss 7 MP microscope (Zeiss Instruments, Thornwood, NY) equipped with a femtosecond-pulsed tunable Ti:Sapphire laser (Coherent, Santa Clara, CA), tuned to a wavelength of 940 nm (<40 mW at back aperture). Fluorescence images were acquired using a 20X/1.0 water immersion objective. GCaMP6s signal was filtered through a green pass band filter (500-550 nm), while red pipette fluorescence was filtered through a 575-610 nm band filter. Fluorescence images were acquired at either low (∼2.5 Hz) or high (>30 Hz) imaging rate for ground truth recordings *ex vivo* (Table 1) and at ∼3 Hz *in vivo*.

**Table 1.** Overview of recorded ground truth datasets. The first data row indicates the recording characteristics for GABAergic neurons, the second data row the recording characteristics for glutamatergicneurons. Recordings with large FOVs and low frame rates (<3 Hz) exhibit higher noise levels compared to zoomed-in FOVs with higher frame rates (>30 Hz), as described in the main text. The total recording duration (third column) is the recording duration for both slow (<3 Hz) and fast (>30 Hz) recordings. The last column (“Spikes within bursts”) quantifies the fraction of spikes with neighboring spikes closer than 10 ms. Values are displayed as mean ± standard deviation, except for spike rate, which is reported as median ± standard deviation for consistency with Fig. 2a. Standardized noise levels were computed as the median absolute fluctuations of ΔF/F, normalized by the imaging frame rate (Methods).

|  | Number of neurons | Total recording duration (min) | Recording duration (<3 Hz) (min) | Noise (<3 Hz) (%·Hz<sup>-1/2</sup>) | Recording duration (>30 Hz) (min) | Noise (>30 Hz) (%·Hz<sup>-1/2</sup>) | Spike rate (Hz) | Spikes within bursts (%) |
| --- | --- | --- | --- | --- | --- | --- | --- | --- |
| GABAergic neurons | 23 | 9.6 ± 3.7 | 4.6 ± 1.6 | 4.3 ± 1.1 | 4.9 ± 1.9 | 1.0 ± 0.3 | 1.1 ± 3.6 | 6.5 ± 8.0 |
| Glutamatergic neurons | 21 | 10.7 ± 6.0 | 5.0 ± 3.3 | 7.2 ± 2.9 | 5.7 ± 2.9 | 1.0 ± 0.2 | 3.7 ± 2.9 | 15 ± 33 |

### Electrophysiological recordings

For ground truth recordings, calcium imaging was performed as described above together with electrophysiological recordings of the same neuron.

Superficial dorsal horn neurons with spontaneous or dorsal root-evoked activity were identified and targeted for cell-attached recording. A pipette microelectrode was pulled from borosilicate glass capillaries (B150F-4, World Precision Instruments, Sarasota, FL) with the Model P-97 micropipette puller (Sutter Instrument, One Digital Drive, Novato, CA) to achieve a tip resistance of ∼4-6 MΩ. The pipette was filled with ACSF containing 30 µM CF594 (Sigma– Aldrich) filtered with a 0.22 µm filter and mounted on an MP-225 motorized micromanipulator (Sutter Instrument). Positive pressure was applied, and the pipette tip was guided into the dorsal horn using the two-photon microscope by switching between the green (for GCaMP) and red (for CF594) channels to approach the target neuron. Then, pressure was released, and a small amount of negative pressure was used to attach to the membrane. Extracellular action potentials were amplified with a differential amplifier (Model 3000, A-M Systems, Sequim, WA). Signals were high-pass filtered at 300 Hz and low-pass at 10 kHz. They were sampled at 10 kHz with Digidata1440A and recorded with Clampex 10.7 (Molecular Device/Axon Instruments, San Jose, CA). The *ex vivo* preparation cannot cover natural responses to external input due to the severed connections to ascending and descending pathways. We therefore used dorsal root stimulation to increase the spectrum of neuronal activity covered by our ground truth recordings. Dorsal root stimulation was delivered, as described previously (Fan et al., 2022), via a tight-fitting, thin-wall glass pipette (1.2 mm diameter; Sutter Instruments, Novato, CA) backfilled with ACSF and attached to the root via suction, typically at L4. Dorsal root stimulation consisted of square pulses (0.2 ms duration) at 1 Hz, 500 µA intensity. Stimulation was delivered using a stimulus isolator (A365, WPI, Sarasota, FL) driven by a waveform generator (Pulsemaster A300, WPI). On average, neurons received 4.2 ± 6.7 stimulations (mean ± s.d. across neurons), corresponding to one stimulation every 135 s on average. GABAergic neurons were stimulated frequently often than glutamatergic neurons (1.8 ± 2.8 stimulations for glutamatergic and 6.4 ± 8.3 for GABAergic neurons), and both datasets included neurons that were not stimulated at all. All stimulation responses were also included to train CASCADE models in order to make CASCADE robust towards such evoked activity patterns. Furthermore, our publicly available ground truth datasets acquired for spinal cord (https://github.com/HelmchenLabSoftware/Cascade) contains not only spike times but also the time points of dorsal root stimulation for each recording.

### *In vivo* lumbar spinal cord calcium imaging

To apply spike rate inference to *in vi*vo data, we used previously acquired calcium imaging datasets that were based on an imaging window above the lumbar spinal cord (modified from (Farrar et al., 2012)) and that targeted the same two classes of neurons as in ground truth recordings (Sullivan and Sdrulla, 2022). All experiments were acquired at ∼3 frames per second with the same two-photon microscope used for *ex vivo* imaging experiments. Anesthetic experiments were carried out by exposing mice to 1% isoflurane for at least 10 minutes prior to 5-minute recordings of activity under this condition. The isoflurane concentration was then increased to 2% and the same incubation and recording periods ensued.

### Extraction of simultaneous calcium imaging and electrophysiology recordings

For calcium imaging movies, recorded neurons were outlined in ImageJ, and the raw fluorescence was averaged for each region of interest. ΔF/F_0_ was calculated with F_0_ being the 10^th^ percentile of raw fluorescence values. For electrophysiological recordings, action potentials were detected using a threshold to extract an initial template for refined matching based on the generated template (Pernía-Andrade et al., 2012), followed by visual inspection of all detected action potentials, as described before (Rupprecht et al., 2021). Electrophysiological and extracted ΔF/F_0_ were aligned using time stamps at the beginning and end of each recording. The extracted ground truth datasets are available online together with existing ground truth datasets in a format that can be accessed both in MATLAB and Python (datasets DS#40 and DS#41 on https://github.com/HelmchenLabSoftware/Cascade).

### Extraction of neuronal traces from *in vivo* recordings

Calcium imaging movies were processed with non-rigid motion correction (Pnevmatikakis and Giovannucci, 2017). Next, neuronal ROIs were extracted manually based on their anatomy and based on their functional response using the map of local correlations with an interactive user interface as described previously (Rupprecht and Friedrich, 2018). From the extracted raw ROI traces, ΔF/F_0_ was computed with the baseline level F_0_ defined as the 10^th^ percentile of fluorescence values. To infer spike rates from ΔF/F_0_ traces, we applied CASCADE models trained on *ex vivo* glutamatergic and GABAergic spinal cord neurons (available as *Spinal_cord_inhibitory_3Hz_smoothing400ms_high_ noise* and *Spinal_cord_excitatory_3Hz_smoothing400ms_high_ noise* via https://github.com/HelmchenLabSoftware/Cascade).

### The mouse cortex dataset

For the ‘cortex dataset’, fluorescence traces were downloaded from https://portal.brain-map.org/explore/circuits/oephys, extracted and processed as described before (Rupprecht et al., 2021). The data consist of neuronal ground truth recordings in cortical layer 2/3 during anesthesia from four transgenic mouse lines that express in specific cortical excitatory subpopulations (Huang et al., 2021). Neuropil correction (neuropil contamination ratio, 0.7) was performed and ΔF/F_0_ was computed using a 6-s running 10^th^ percentile window (adjusted to the noisiness of each recording). Importantly, ground truth recordings were carefully quality-controlled as described (Rupprecht et al., 2021).

### Extraction of the calcium response for the average action potential

The calcium response for the average action potential for each neuron (*i*.*e*., the linear kernel; Fig. 3a) was extracted by regularized deconvolution using the *deconvreg(calcium,spikes)* function in MATLAB (MathWorks). This function computes the kernel, which, when convolved with the observed *spikes*, results in the best approximation of the *calcium* trace. The same linear kernel was used for the linear forward model to predict the linear expected response (Fig. 3e).

### Computation of noise levels

Standardized noise levels were computed, as described previously in more detail (Rupprecht et al., 2021), as the median absolute fluctuations of *ΔF/F* between adjacent timepoints, normalized by the square root of the imaging frame rate *f*_*r*_:

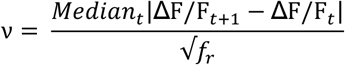

When computed for ΔF/F data, *ν* is quantitatively comparable across datasets, even when frame rates differ, hence the name “standardized noise levels”. The units for *ν* are %·Hz^−1/2^, which we omit in the main text for readability.

### Quantification of spike rate inference performance

Ground truth spike rates used for training and evaluation were generated from discrete ground truth spikes by convolution with a Gaussian smoothing kernel. To take advantage for the respective sampling rate, the precision of the ground truth was adjusted by tuning the standard deviation of the smoothing Gaussian to the temporal sampling rate (σ = 0.4 s for 2.5-Hz recordings, σ = 0.2 s for 5 Hz, σ = 0.1 s for 10 Hz, σ = 0.1 s for 15 Hz, σ = 0.1 s for 20 Hz, σ = 0.05 s for 25 Hz and σ = 0.05 s for 30 Hz recordings unless otherwise stated). This smoothed ground truth spike rate was then compared to the inferred spike rate.

To generate ground truth at specific frame rates and noise levels from the existing ground truth, ΔF/F traces were temporally resampled and Gaussian noise was added until the desired noise level was reached, as described previously (Rupprecht et al., 2021). Therefore, for a given condition (e.g., standardized noise level of “7” and sampling rate of 30 Hz), ground truth from a neuron recorded at a standardized noise level larger than the desired level or at an imaging rate slower than the desired sampling rate was used. This procedure enabled the consistent evaluation of algorithms across datasets and individual neurons for standardized ground truth at specific frame rates and noise levels.

### Spike rate inference with CASCADE and OASIS

For spike rate inference with OASIS, the Python implementation of the algorithm in Suite2p (Pachitariu et al., 2019) was downloaded from https://github.com/MouseLand/suite2p and used in Python 3.9 with default parameters. For evaluation of spike rate inference results with OASIS, we tested Gaussian smoothing kernels of variable standard deviation and temporal shifts between −1 and +1 s to find the amount of smoothing and the delay for each dataset that optimized the correlation with ground truth. For spike rate inference with “default CASCADE”, pretrained models together with the algorithm were downloaded from https://github.com/HelmchenLabSoftware/Cascade.

These models had been trained on a large database of excitatory neurons across different brain areas but focused on cortical recordings (“global models” in CASCADE). Within this study, these models are called “default CASCADE” because they were not retrained with ground truth acquired from spinal cord.

### Retraining CASCADE for spike rate inference with spinal cord neurons

CASCADE was retrained from scratch on spinal cord ground truth with the same procedures as described before (Rupprecht et al., 2021). The retrained CASCADE networks consist of a standard convolutional network with six hidden layers, including three convolutional layers. The input consists of a window of 64 time points (32 time points for frame rates <15 Hz), symmetric around the time point for which the inference was made. The three convolutional layers have relatively large filter sizes (31, 19 and 5 time points; 17, 9 and 3 time points for frame rates <15 Hz), with an increasing number of features (20, 30 and 40 filters per layer), with max pooling layers after the second and third layer, and a densely connected hidden layer consisting of ten neurons as the final layer. Models for spinal cord neurons were trained separately on glutamatergic and GABAergic neurons, including all available neurons, except for GABAergic neurons with extremely high outlier spike rates that would otherwise strongly bias the training. Care was taken not to use the same neuron for training and testing of a model. For example, if a model trained at a frame rate of 30 Hz with the glutamatergic dataset was tested on glutamatergic neurons, a new model was trained for each test neuron on all glutamatergic spinal cord neurons except the neuron tested for (“leave-one-out” strategy). This strategy of cross-validation prevents fitting of test data and enables us to test the generalization of the algorithm to unseen data. Models retrained with spinal cord data are already integrated into CASCADE (https://github.com/HelmchenLabSoftware/Cascade) for a subset of sampling rates (2.5 Hz, 3 Hz and 30 Hz); further models tailored towards special use cases can be readily requested as described on this Github page.

### High-frequency spike events

For the analysis of short events with a large number of spikes (“high-frequency spike events”; Fig. 5), we used the selection criterion that the maximum instantaneous spike rate of the ground truth exceeded 45 Hz. Changes to the threshold resulted in qualitatively similar results.

### Experimental design and statistical analysis

The number of recorded ground truth neurons (approx. 20 per group) was similar to previous studies that analyzed such ground truth recordings (Tian et al., 2009; Chen et al., 2013; Huang et al., 2021). The animal’s sex was not included as analysis variable due to the limited sample size. Statistical analyses were performed in MATLAB 2020a. Only non-parametric tests were used. The Mann–Whitney rank-sum test was used for non-paired samples, and the Wilcoxon signed-rank test was used for paired samples. Two-sided tests were applied unless otherwise stated. Box plots used standard settings in MATLAB, with the central line at the median of the distribution, the box at the 25th and 75th percentiles and the whiskers at extreme values excluding outliers (outliers defined as data points that are more than 1.5·D away from the 25th or 75th percentile value, with D being the distance between the 25th and 75th percentiles).

### Software Accessibility

CASCADE software, together with CASCADE models trained on glutamatergic or GABAergic spinal cord ground truth and the ground truth datasets are available online via https://github.com/HelmchenLabSoftware/Cascade.

## Results

### Ground truth recordings for calcium imaging in mouse spinal cord

To understand how calcium signals in spinal cord neurons relate to the underlying action potentials and how this relationship might differ between excitatory and inhibitory cells, we performed simultaneous electrophysiological recordings and calcium imaging in the intact, *ex vivo* spinal cord isolated from transgenic mice expressing GCaMP6s either in vGluT2-(glutamatergic) or vIAAT-positive (GABAergic) cells (Fan et al., 2022) (Fig. 1a,b; Methods). Cell-attached electrical recordings were established on individual GCaMP6s-expressing neurons, as identified under the two-photon microscope, permitting the simultaneous recording of action potentials (spikes) and calcium signals from the same cell (Fig. 1c-d). We recorded from 21 glutamatergic and 23 GABAergic neurons and collected ground truth data with >70 000 action potentials over a total recording duration of 7.4 hours. Recordings were performed in two imaging configurations, either acquiring movies with a field-of-view (FOV) used for population imaging (512^2^ pixels; approx. 425 µm side length) at a low frame rate of approximately 2.5 Hz (Fig. 1c,d), or with a smaller imaging FOV (35^2^ −45^2^ pixel, approx. 30-35 µm side length) at a higher frame rate (33 ± 8 Hz, mean ± s.d.; inset in Fig. 1c,d). The simultaneously performed electrophysiological recordings exhibited a broad diversity of spiking and bursting patterns, with variable spike rates (SR) between as well as within excitatory and inhibitory cell types (Fig. 1e-j). More details about the summary recording statistics are provided in Table 1.

**Figure 1.**
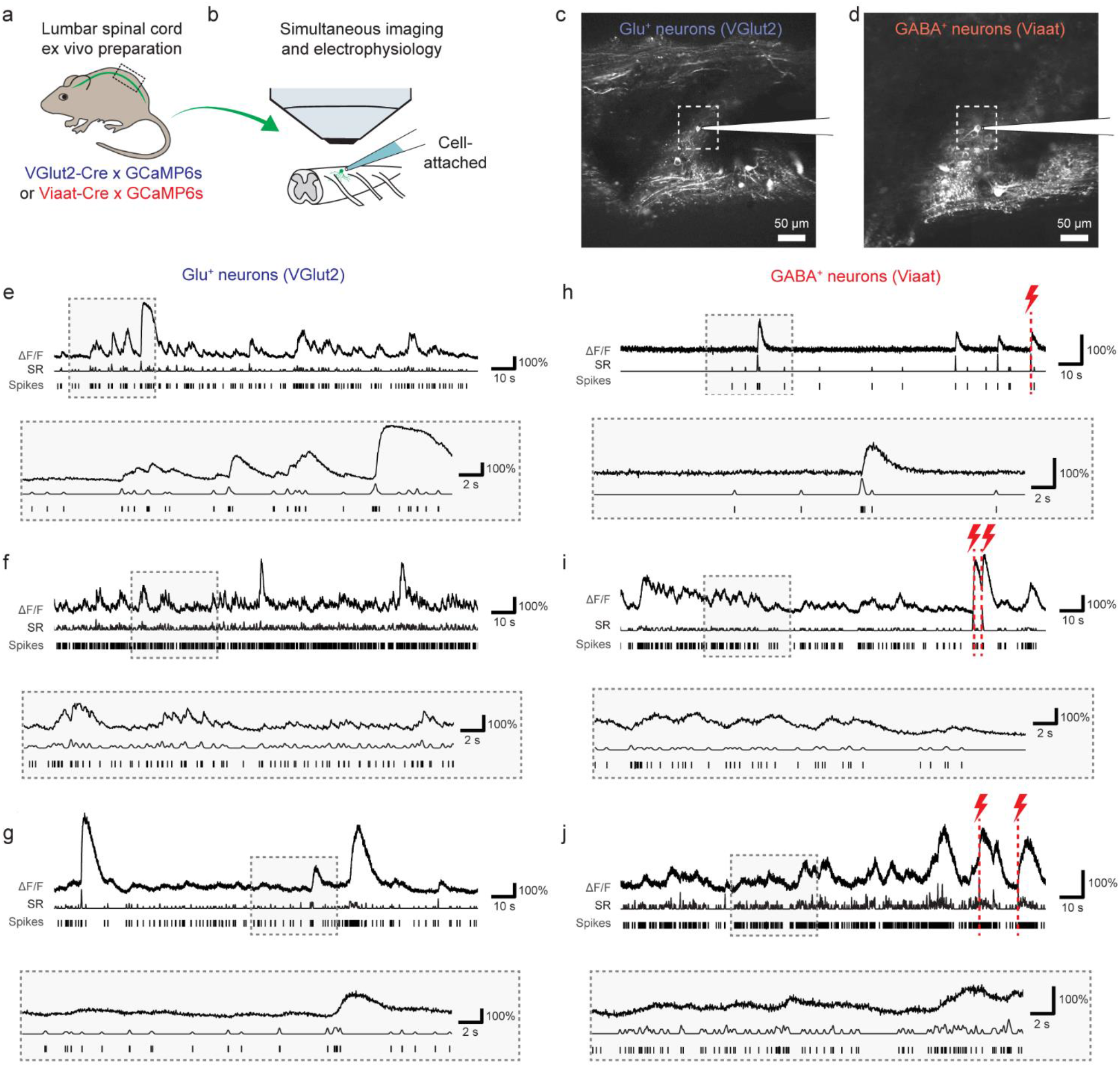
Simultaneous recording of electrophysiological spikes and calcium signals in glutamatergic and GABAergic neurons of mouse spinal cord. **a**, Scheme of transgenic expression of GCaMP6s in glutamatergic (blue) and GABAergic neurons (red) in mouse spinal cord. **b**, Lumbar spinal cord explant preparation used for ground truth recordings with simultaneous two-photon calcium imaging and cell-attached recordings. **c**, Example FOV for a slow recording (2.5 Hz) for glutamatergic neurons (VGlut2). The sub-area highlighted by the dashed rectangle indicates the FOV for a faster recording (>30 Hz) from the same neuron. **d**, Same as in (c) but for a GABAergic neuron (Viaat). **e-g**, Examples of ground truth recordings from three glutamatergic neurons, with zoom-in to a subregion below. For each recording, the normalized fluorescence extracted from calcium imaging (ΔF/F), the smoothed spike rate derived from electrophysiological spike times (SR), and the spike times detected from the electrophysiological recording (Spikes) are shown. **h-j**, Same as in (e-g) but for three example GABAergic neurons. Red flashes indicate times when dorsal root stimulation was applied.

### Variability of spike patterns across neurons and datasets

First, we systematically analyzed the spike patterns for glutamatergic and GABAergic spinal cord neurons from the ground truth recordings and compared the results with previously obtained ground truth datasets from GCaMP-expressing mouse cortical neurons, hence called the ‘cortex dataset’ (Huang et al., 2021).

This cortex dataset consists of ground truth recordings from 80 neurons in animals with four different transgenic strategies to express either GCaMP6s or GCaMP6f in layer 2/3 pyramidal cells of visual cortex (Fig. 1-1 for examples of recordings) (Huang et al., 2021).

First, we systematically analyzed the spike patterns for glutamatergic and GABAergic spinal cord neurons from the ground truth recordings and compared the results with previously obtained ground truth datasets from GCaMP-expressing mouse cortical neurons, hence called the ‘cortex dataset’ (Huang et al., 2021). This cortex dataset consists of ground truth recordings from 80 neurons in animals with four different transgenic strategies to express either GCaMP6s or GCaMP6f in layer 2/3 pyramidal cells of visual cortex (Fig. 1-1 for examples of recordings) (Huang et al., 2021).

For spinal cord ground truth recordings, we observed that spike rates varied by an order of magnitude within both the glutamatergic and the GABAergic spinal cord dataset (Fig. 2a). Spike rates of glutamatergic neurons in these recordings were, on average, significantly higher compared to GABAergic neurons (p = 0.001, Wilcoxon rank sum test) and distributed more uniformly across a broader range (Fig. 2a; but notice the outlier GABAergic neuron with very high spike rate).

**Figure 2.**
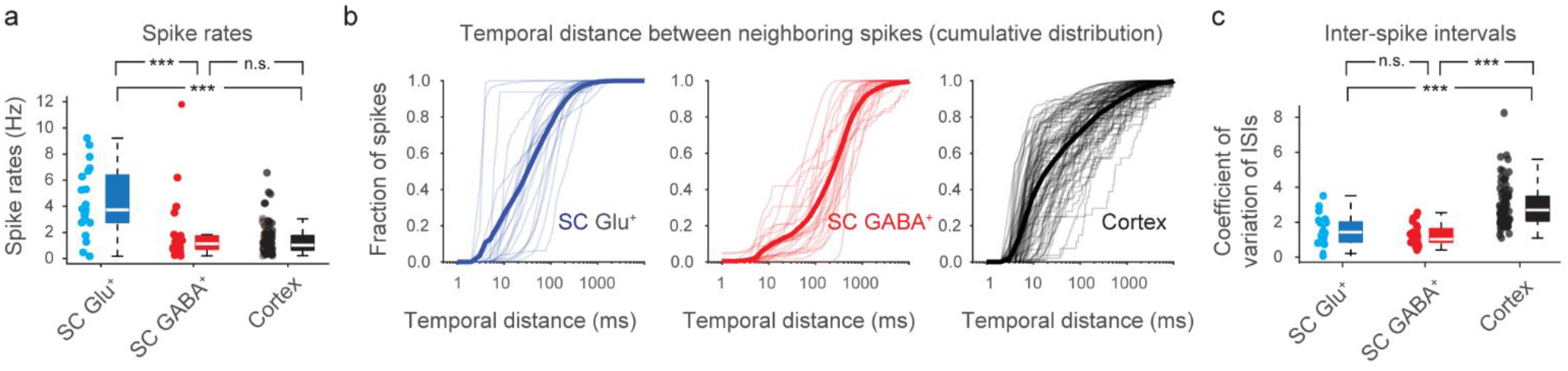
Comparison of electrophysiological characteristics of ground truth recordings from spinal cord and cortex. **a**, Spike rates for spinal cord glutamatergic neurons ground truth (*SC Glu*^*+*^, blue), spinal cord GABAergic neurons ground truth (*SC GABA*^*+*^, red), both recorded for this study with GCaMP6s, and excitatory neurons in transgenic mouse cortex that express GCaMP6s or GCaMP6f (*Cortex*, black). Each data point underlying the box plot represents a recorded neuron (n = 21, 23 and 80 neurons for *SC Glu*^*+*^, *SC GABA*^*+*^ and *Cortex*). Spike rates of *SC Glu*^*+*^ were higher than for *SC GABA*^*+*^ and *Cortex* (p = 0.001 and 0.000001), with no significant difference between the latter two (p = 0.83, Wilcoxon rank sum test). **b**, Cumulative distribution of the temporal distance from each spike to the neighboring spike, as a measure of burstiness. Spinal cord spike patterns exhibit lower burstiness compared to spike patterns from the cortex dataset. Each thin line indicates the cumulative distribution of these distances from all spikes of an individual neuron. Bold lines indicate the average across neurons within a dataset. **c**, Coefficient of variation (COV) of inter-spike intervals (ISIs) as an additional measure of burstiness. Higher COV indicates higher burstiness, lower COV a less bursty and more regular spike pattern. As in (a), each datapoint is from an individual neuron. COV was significantly higher for the cortex dataset, reflecting higher burstiness (p < 10^−6^, Wilcoxon ranksum test); no difference between spinal cord datasets was found (p = 0.24). *p < 0.05, **p < 0.01, ***p < 0.001, n.s., not significant (p > 0.05).

The relationship between neuronal spiking and the observed fluorescence signal from calcium indicators is typically non-linear. For example, it is possible that a neuron exhibits no discernible calcium response to a single spike but a strong calcium response to two or three closely spaced spikes. Therefore, the electrophysiological spike patterns of a neuron will influence how well information about spikes can be extracted from calcium imaging data. We thus investigated the bursting propensity of the recorded spinal neurons and compared it to existing cortical datasets. To this end, we went through all spikes recorded for a given neuron and measured the temporal distance between each spike and its closest neighboring spike. This procedure is similar to the computation of inter-spike intervals but searches for the closest neighboring spike in either the past or future and therefore measures more accurately whether a given spike is part of a burst or not. Then, we plotted the cumulative distribution of closest temporal neighbors for each neuron (Fig. 2b). Based on this analysis, we found that the bursting propensity (defined here as the fraction of spikes that were within 10 ms of any neighbouring spike) was highest for the neurons in the cortex dataset (42 ± 23%, median ± s.d. across neurons), followed by the glutamatergic (16 ± 33%) and the GABAergic (6.5 ± 8.0%) spinal cord neuron datasets. This result was supported by a complimentary analysis measuring the coefficient of variation (COV) of inter-spike intervals as a standard measure of burstiness (Softky and Koch, 1993) (Fig. 2c), displaying a marked increase of the COV for the cortex dataset compared to the spinal cord datasets (p < 10^−6^) but no difference between the excitatory and inhibitory spinal cord datasets (p = 0.24). These analyses reveal that glutamatergic and particularly GABAergic neurons recorded in our spinal cord ground truth exhibited less bursty firing patterns than neurons from the cortex dataset but rather fired more steadily and regularly as indicated by the COV. These differences further emphasize the issue of how well algorithms for spike rate inference generalize from cortex to spinal cord data.

### Magnitude and variability of spike-evoked calcium signals

Next, we examined the variability of the evoked calcium signals in response to a well-defined electrophysiological event (Fig. 3a-c). To this end, we took advantage of the fact that calcium signals can be approximated as a convolution of spikes with a kernel function (*i*.*e*., the calcium response evoked by the average spike). The kernel can be easily retrieved from the ground truth by deconvolution of the recorded calcium signals (ΔF/F) with the simultaneously recorded spikes, as shown before (Rupprecht et al., 2021). The kernel therefore constitutes the ΔF/F response not to isolated spikes but to the average spike. We found that the resulting calcium response kernels for both glutamatergic and GABAergic spinal cord neurons exhibited visibly slower time courses compared to previously performed recordings in mouse cortex with GCaMP6s and GCaMP6f (Huang et al., 2021) (single-exponential fits for the time constant τ yielded 3.1 ± 0.5 s, fit ± 90% confidence interval, for glutamatergic spinal cord neurons; 4.4 ± 0.7 s for GABAergic neurons; 0.8 ± 0.1 s for the cortex dataset). In a subset of recordings, we performed physiological (37°C) instead of room temperature recordings but did not observe a visibly faster calcium response function (Fig. 2-1; τ = 2.9 ± 0.5 s for room temperature, 3.0 ± 0.6 s for physiological temperature). For the response kernel amplitudes as measured by peak response (maximum ΔF/F) and mean response (average ΔF/F during the 2 s post-event window), we observed a striking variability on a neuron-to-neuron basis for spinal cord datasets (Fig. 3b,c; blue and red) and the mouse cortex dataset (Fig. 3b,c; black). This is consistent with the previously found variability of response amplitudes from neuron to neuron (Éltes et al., 2019; Rupprecht et al., 2021). The response magnitude was consistent with previous similar analyses (Rupprecht et al., 2021). We wondered whether the response kernel could be distorted by the inclusion of dorsal root stimulations, because such stimulations might recruit slowly acting neuromodulators (Geppetti et al., 2015; Todd and Spike, 1993) that may influence intracellular calcium (Warwick et al., 2022); alternatively, stimulation may lead to bursts and prolonged membrane depolarizations in the recorded neuron, events that have been associated with prominent calcium signals (Ledochowitsch et al., 2020; Milicevic et al., 2024). However, we did not observe a striking difference when including the stimulation periods (average response as in Fig. 3b, median ± s.d., 12 ± 6% and 11 ± 7%, for glutamatergic and GABAergic neurons) and when excluding the stimulation periods (12 ± 7% and 11 ± 8%), indicating that dorsal root stimulation did not elicit responses distinct from spontaneous activity.

**Figure 3.**
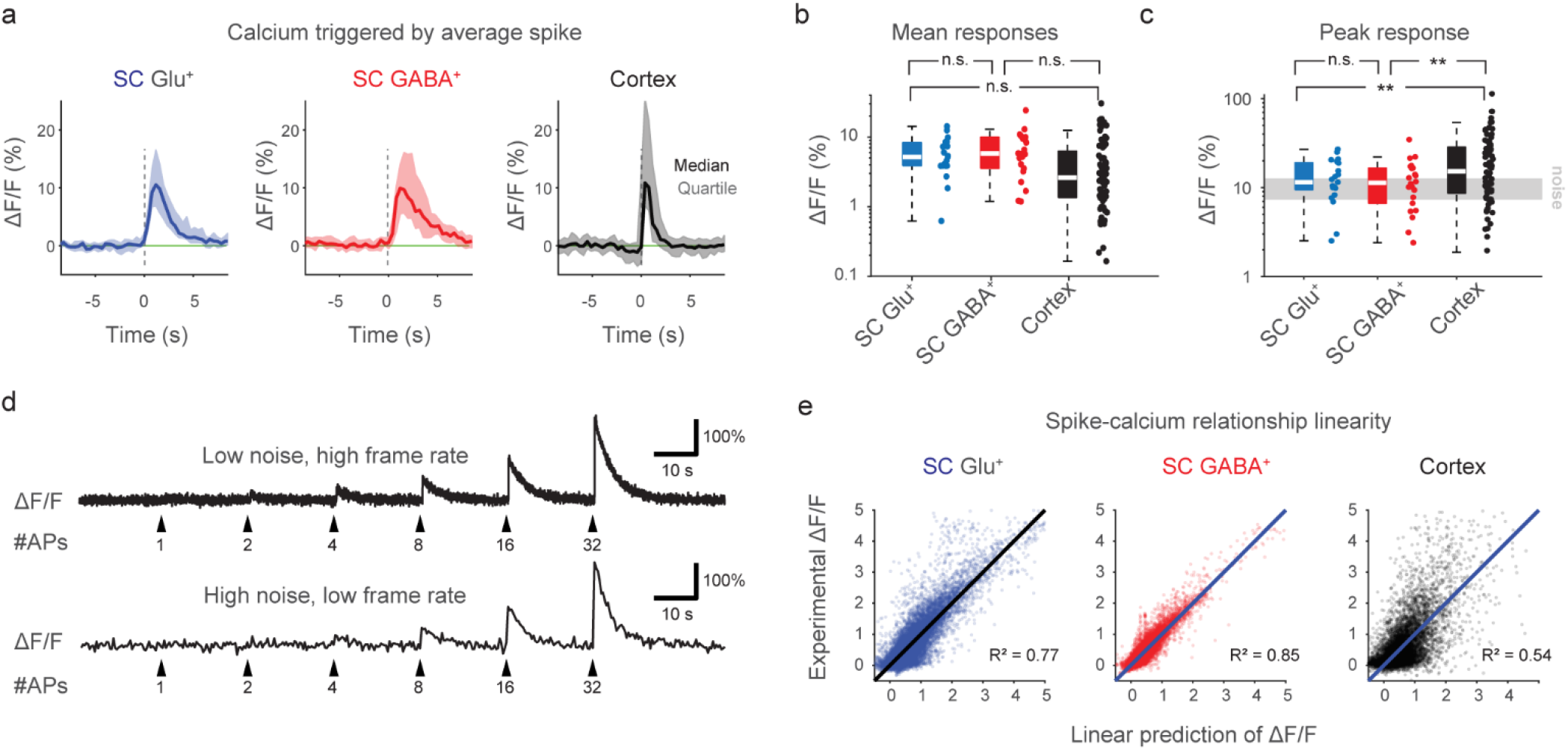
Comparison of the spike-calcium relationship of ground truth recordings from spinal cord and cortex. **a**, Calcium response (ΔF/F) for the average action potential across neurons, computed by linear deconvolution of the ground truth recording, at the common sampling rate of 2.5 Hz. Note that this kernel was derived from all spikes, not only from isolated spikes. Shown is median and quartile corridors across all neurons (n = 21, 23 and 80 for the glutamatergic, GABAergic and cortex datasets). **b**, Mean calcium response (ΔF/F) during the first 2 s after t = 0 s for the average action potential across neurons. Please note the logarithmic y-scale. No significant difference between *SC Glu*^*+*^ and *SC GABA*^*+*^ (p = 0.69) *SC Glu*^*+*^ and *Cortex (p = 0*.*17)* or *SC GABA*^*+*^ and *Cortex* (p = 0.06, Wilcoxon rank sum test). **c**, Peak calcium response (ΔF/F) for the average action potential across neurons. Please note the logarithmic y-scale. No significant difference between *SC Glu*^*+*^ and *SC GABA*^*+*^ (p = 0.52), but increase of *Cortex* compared to *SC Glu*^*+*^ *(p = 0*.*0071)* or *SC GABA*^*+*^ (p = 0.0049, Wilcoxon rank sum test). **d**, Illustrative simulation of ΔF/F traces with typical noise levels and calcium transients evoked by average (median) amplitude (SC Glu^+^ kernel in panel (a)). The simulated recording conditions are 40 Hz imaging rate and a standardized noise level (Methods) of “1.0” for the upper trace, and 2.54 Hz and noise level of “5.0” for the lower trace. Spike-evoked calcium transients can be clearly observed for a large number of simultaneous spikes but not distinguished from noise for single spikes. **e**, The experimental ΔF/F values are predicted by the convolution of spike times with the linear kernel (from panel (a)). The plot indicates how well a neuron-specific linear forward model fits the experimental data. Each visualized data point is a timepoint from a ground truth recording downsampled to 3 Hz (number of timepoints: 27’816 for Glu^+^, 28’835 for GABA^+^, 31’613 for cortex). *p < 0.05, **p < 0.01, ***p < 0.001, n.s., not significant (p > 0.05).

The average peak response of a subset of neurons surpassed the typical levels of ΔF/F fluctuations, but for the large fraction of neurons this was not the case (Fig. 3c). From a simulation based on linear convolution of calcium kernels with realistic noise levels (Fig. 3d), calcium responses cannot be visually discerned for <2-4 quasi-simultaneous spikes for low-noise recordings (standardized noise level of 1.0, as defined in Methods, frame rate of 40 Hz) and for <4 spikes for high-noise recordings (noise level of 5.0, frame rate of 2.5 Hz). The real response to isolated single spikes might be even smaller due to the known sigmoid non-linearity of GCaMP6 (Rose et al., 2014).

Finally, we attempted to visualize the linearity of the calcium indicator responses to spike rates for each dataset. In principle, the same indicators should yield a similar calcium vs. fluorescence relationship across cells, but only if the respective intracellular calcium ranges are comparable. However, due to the variability of typical calcium concentrations across cell types (Maravall et al., 2000) and the sigmoid calcium concentration response curve of GCaMPs (Rose et al., 2014), the same indicator might behave linear in one condition, supra-linear in another and saturating in yet another. A typical method to quantify linearity is to measure responses evoked by a set of one, two or more isolated action potentials (Kerr et al., 2005; Lütcke et al., 2010). However, in our recorded spinal cord ground truth dataset, isolated singlets, doublets or triplets of action potentials occurred only in few neurons, therefore disabling this approach. Instead, we computed the expected fluorescence response from the measured spike pattern as estimated with a linear spike-to-calcium forward model (Wei et al., 2020). In this forward model, we employed linear convolution of the spike pattern with the calcium response kernel as computed before (Fig. 3a). We observed that the experimentally measured ΔF/F was reasonably linear according to such a forward model for GABAergic spinal cord neurons and, to a lesser extent, for glutamatergic spinal cord neurons (Fig. 3e). In contrast, such a linear relationship was less obvious in the cortex dataset (Fig. 3e). This qualitative finding shows that the experimentally measured ΔF/F reflects changes of the gradually and slowly changing underlying spike rates in most of the ground truth recordings in spinal cord (see also Fig. 1e,f,g,I,j). This result is reminiscent of observations made for cortical interneurons (Inoue et al., 2019; Kwan and Dan, 2012) but in contrast to the cortex datasets from excitatory neurons, in which isolated calcium transients due to bursts (Fig. 2b,c) rather than regular firing were typically observed (Chen et al., 2013; Huang et al., 2021). This finding of gradually and relatively linearly changing fluorescence suggests that raw ΔF/F might be considered as a useful approximation of spike rates for spinal cord neurons without any spike inference algorithm applied.

### Spike rate inference from spinal cord calcium imaging data

Next, we wanted to benchmark methods to infer spiking activity from calcium imaging data in spinal cord. To address this challenge, we used several complementary approaches (Fig. 4a). First, following up on our investigation of calcium response linearity, we used the raw ΔF/F signal as a proxy for spiking activity. Second, we used OASIS, an unsupervised spike rate inference algorithm based on non-negative deconvolution that is used as default option in the two most widely employed toolboxes for source extraction of calcium imaging data (CaImAn, Suite2p) (Giovannucci et al., 2019; Pachitariu et al., 2019). Third, we used a supervised spike rate inference algorithm (CASCADE) trained on a diverse and large dataset that mostly contained recordings from mouse cortex and no neurons from spinal cord (Rupprecht et al., 2021). Finally, we retrained CASCADE on our specifically recorded ground truth spinal cord datasets containing either glutamatergic or GABAergic neurons. Among these algorithms, only the CASCADE models provide an estimate of absolute spike rates. Importantly, to test these retrained supervised CASCADE models on our spinal data sets, we applied a “leave-one-out” principle, where CASCADE was retrained each time with the “test neuron” excluded. This procedure enabled us to test the generalization of the algorithm without reducing the dataset. To standardize the benchmarking, we used experimentally recorded datasets but for each recording added Gaussian noise until a certain ‘standardized noise level’ (e.g., a noise level of “7”) (Methods) was reached. Similarly, we used downsampling of recorded data to generate a benchmarking dataset of a specified frame rate (e.g., 30 Hz). This procedure, as introduced previously (Rupprecht et al., 2021), standardizes performance evaluations and makes them comparable across neurons within datasets and across datasets.

**Figure 4.**
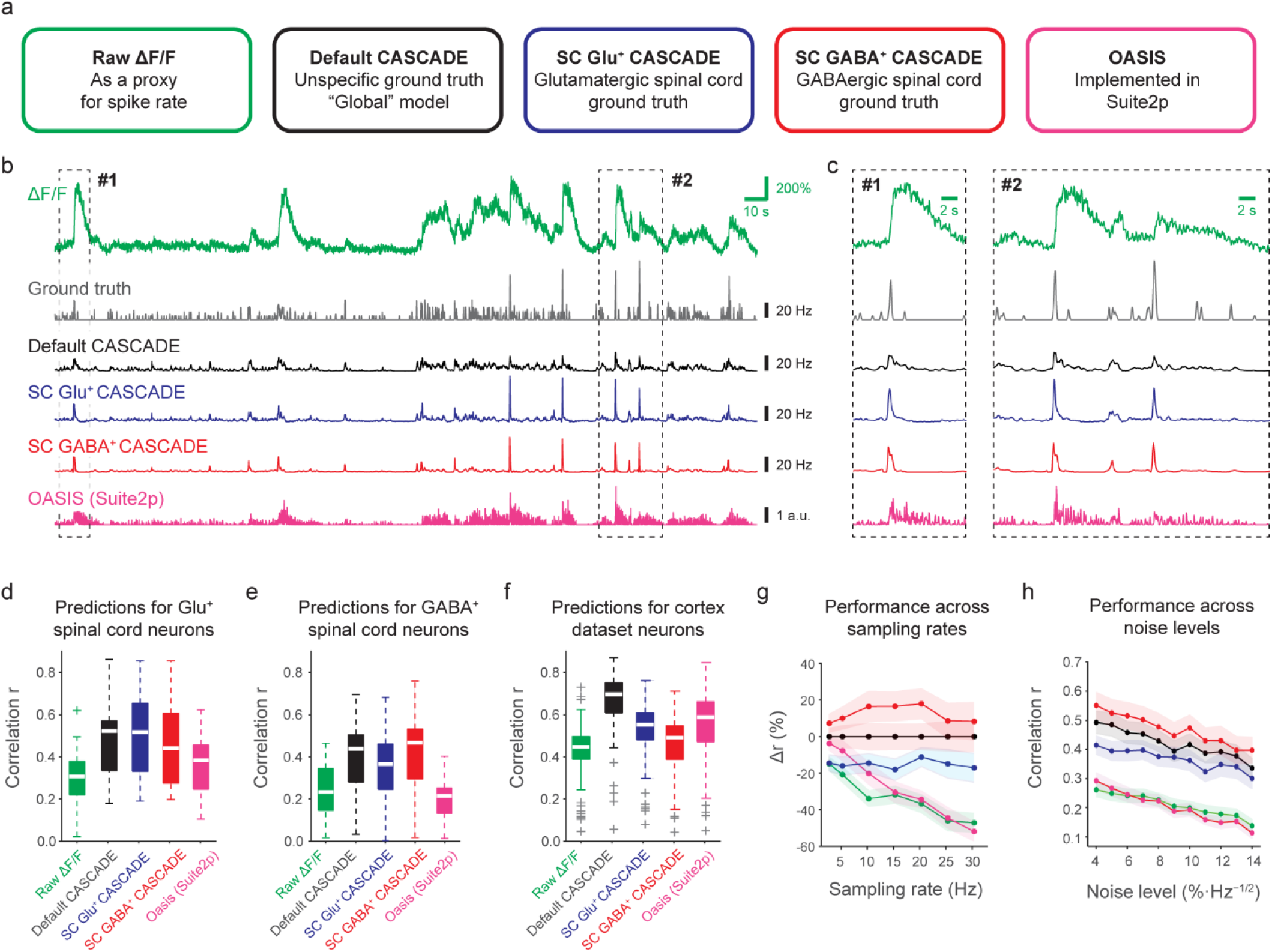
Spike rate inference algorithms in spinal cord neurons. **a**, Schematic of five different approaches that can be used as a proxy readout for neuronal spike rates: (1) Raw ΔF/F (green), (2) Spike rate inference with CASCADE, trained on a diverse ground truth dataset that mostly consists of cortical excitatory neurons (black), (3) Spike rate inference with CASCADE, trained on a ground truth dataset with glutamatergic spinal cord neurons (blue), (4) Spike rate inference with CASCADE, trained on a ground truth dataset with GABAergic spinal cord neurons (red), (5) Unsupervised spike rate inference using OASIS, implemented in Suite2p (magenta). **b**, Examples of an extracted ground truth recording together with the ground truth spike rate and spike rate predictions with default and retrained CASCADE, as well as OASIS. Sampled at 30 Hz with a standardized noise level of “7”, from an excitatory neuron. The same ground truth with spike rate predictions resampled at 2.5 Hz are shown in Fig. 4-1. **c**, Zoom-in to sub-regions in (b), highlighting the differences of predictions for high-frequency spike events. **d-f**, Quantification of performance of the five approaches described in (a) for the glutamatergic spinal cord dataset (d) and the GABAergic spinal cord dataset (e) and the cortex dataset (f). Each datapoint underlying the boxplots is a ground truth recording from a different neuron (n = 21, 23 and 80 for glutamatergic, GABAergic and cortex datasets). Ground truth was resampled at 30 Hz at a standardized noise level of “7” (see Methods). Performance was evaluated after correction for systematic delays (Fig. 4-2). Outcomes of relevant statistical tests are reported in the main text. **g**, Quantification of performance for the GABAergic spinal cord dataset across sampling rates, normalized for the performance of the default CASCADE model. The same quantification for the glutamatergic spinal cord dataset is shown in Fig. 4-3a. Color code as in (a-d). **h**, Quantification of performance for the GABAergic spinal cord dataset across noise levels. The same quantification for the glutamatergic spinal cord dataset is shown in Fig. 4-3b.

To compare these approaches, we performed spike rate inference for all recorded neurons in our two ground truth datasets. As an initial result of our comparison, all algorithms performed reasonably well and yielded meaningful predictions (Fig. 4b-c; Fig. 4-1).

To quantify the performance of each approach, we computed the correlation between predicted and true spike rates as done previously (Berens et al., 2018; Rupprecht et al., 2021; Theis et al., 2016). Such a performance metric, however, puts the ΔF/F approach at a disadvantage since ΔF/F will always be delayed with respect to the spiking ground truth, thereby reducing the correlation with ground truth. A similar systematic delay was also found for model-based spike rate inference algorithms like OASIS in a previous study (Rupprecht et al., 2021). We, therefore, shifted the predictions temporally to optimize the correlation with the same shift applied to all neurons of a given dataset (Fig. 4-2) and used this optimal temporal shift to evaluate a given dataset and approach to estimate spiking activity.

We quantified performance using a fixed sampling frequency (30 Hz) and a fixed noise level of the calcium imaging data. For these fixed settings, typical correlation values as a performance readout (median value across all spinal cord neurons) were ∼0.25 ΔF/F, ∼0.30 for OASIS and ∼0.47 for default CASCADE. Predictions using default CASCADE were overall better than predictions by OASIS or when using ΔF/F as a direct proxy, and this finding held true both for the glutamatergic and GABAergic dataset (Fig. 4d,e; median correlation of 0.p < 0.001 for all comparisons; Wilcoxon signed-rank test; n = 23/21 neurons for the glutamatergic and GABAergic dataset). In addition, we found that a version of CASCADE that was retrained with a specific dataset (sticking to the “leave-one-out” principle) performed equally or slightly better than the default CASCADE algorithm (p-value p = 0.37, increase of median performance Δr = −3.4% for the glutamatergic dataset; p = 0.001, Δr = +8.3% for the GABAergic dataset), and slightly better than the same algorithm trained with the other spinal cord dataset (p = 0.0057, Δr = +9.1%, and p = 0.0006, Δr = +23.4% for glutamatergic and GABAergic dataset, respectively). In addition, we applied all models to the mouse cortex dataset (Fig. 4f) and found that models designed for cortex performed best with default CASCADE, significantly better than all other tested approaches (p < 10^−10^, signed-rank test, n = 80 neurons), followed by OASIS that performed better than CASCADE trained on GABAergic (p < 10^−10^) but not significantly better than CASCADE trained on glutamatergic ground truth (p > 0.05). In addition, we tested whether performance improvements through retraining of CASCADE were stronger for potential subtypes of spinal cord neurons as identified for example by mean firing rates or burstiness, but we did not find such a relationship (Fig. 4-4). These analyses show that models not specifically trained for the new datasets generalize reasonably well between cortical and spinal cord data and that retraining improved the performance, in particular for the GABAergic spinal cord neurons.

These model performances were confirmed across imaging conditions when re-sampling the ground truth dataset at different imaging rates (Fig. 4g; Fig. 4-3a) and across different resampled noise levels (Fig. 4h; Fig. 4-3b). Notably, the performances of all approaches, including the use of raw ΔF/F, although still distinct from each other, were getting closer for very low imaging rates (2.5 Hz; Fig. 4g and Fig. 4-3a). It should be noted that lower imaging rates were accompanied by lower precision of temporal evaluation (*e*.*g*., 400-ms standard deviation Gaussian smoothing for 2.5 Hz imaging rate, and 50-ms standard deviation Gaussian smoothing for 30 Hz imaging rate). As temporal precision was decreased, the performance of the ΔF/F approach for 30-Hz image rate again came closer to the performance of supervised CASCADE (Fig. 4-3c).

Together, these results show that spike rate inference with the supervised CASCADE retrained on a specific dataset outperforms the non-supervised approaches (raw ΔF/F or OASIS). Only for low calcium imaging rates (<< 5 Hz), the performance of the approach using raw ΔF/F or OASIS was almost equally good as the retrained supervised algorithm. Furthermore, a default CASCADE algorithm trained on a diverse dataset that does not include spinal cord neurons already performed excellently and could also be used as a reliable spike rate inference algorithm for both glutamatergic and GABAergic neurons in the spinal cord.

### Reduced spike rate inference performance on high-frequency spike events

Next, we evaluated the performance of the spike rate inference algorithms across different spike patterns. This evaluation was performed at fixed conditions across neurons and datasets (resampled frame rate of 30 Hz, standardized noise level of 7). From an inspection of predicted spike patterns (Fig. 4c), we noticed that events with a large number of spikes in a small time window, hence called high-frequency spike events, were not well recovered by the algorithms designed for or trained with cortical excitatory neurons, *i*.*e*., OASIS or default CASCADE. Specifically, the recovered spike pattern was considerably prolonged compared to the ground truth (Fig. 4c). To quantify this observation, we selected all events where spike rates exceeded a predefined threshold (instantaneous firing rate of 45 Hz, see Methods) (Fig. 5a,f) and plotted the associated ΔF/F values (Fig. 5b,g), the spike rates inferred by default CASCADE (Fig. 5c,h), by retrained CASCADE (Fig. 5d,i) and by OASIS (Fig. 5e,j). We noticed that spike rates inferred by default CASCADE and even more by OASIS exhibited false positive spike detections when the true spike rate had already decayed (Fig. 5a-j; Fig. 5-1). For retrained CASCADE, the inferred spike rate more accurately reflected the ground truth spike rate dynamics around high-frequency spike events (Fig. 5f,I; Fig. 5-1), demonstrating how spike rate inference for spinal cord neurons can be optimized by retraining.

**Figure 5.**
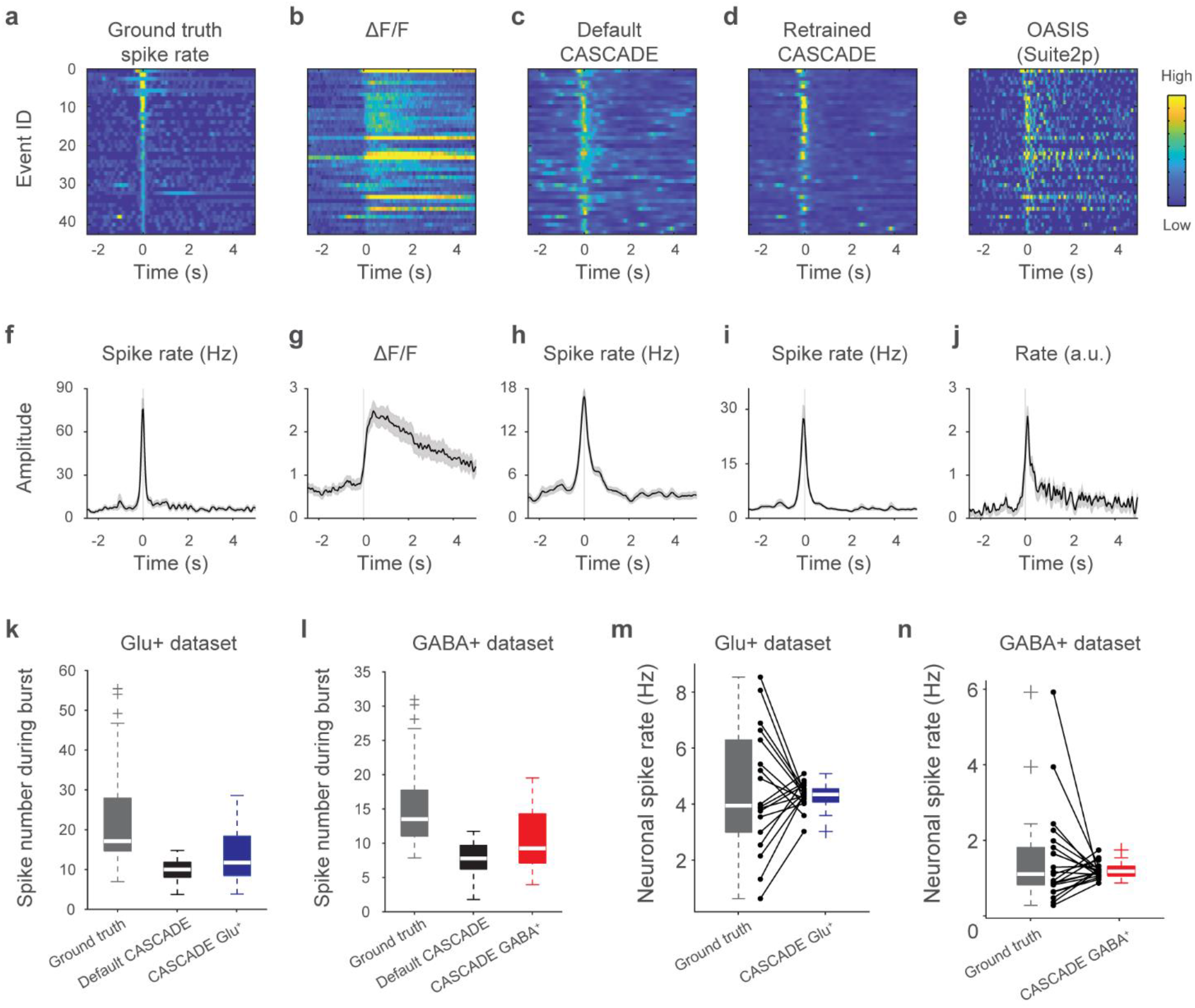
Spike rate inference for high-frequency spike events for glutamatergic neurons in spinal cord. **a-e**, High-frequency spike events for the glutamatergic neuron dataset, with the corresponding associated ground truth spike rate (a), ΔF/F signal (b), spike rate inferred by the default CASCADE model (c), spike rate inferred from the retrained CASCADE model (SC Glu^+^ CASCADE) (d), and spike rate inferred by the OASIS algorithm (e). **f-j**, Same as in (a-e) but averaged across events, with absolute values indicated if possible. Prolonged spike rate is seen for the default CASCADE and OASIS models. The median number of spikes during events (1-s window around event) for ground truth vs. default CASCADE vs. retrained CASCADE is 13.6 vs. 7.9 vs. 9.4 spikes.

Next, we quantified the accuracy of spike rate inference during these high-frequency spike events. For this comparison, OASIS was not included since the OASIS algorithm is not intended to recover absolute spike rates. We found that spike rates in a 1-s window around high-frequency spike events were systematically underestimated by all versions of CASCADE, probably due to indicator saturation, for both spinal cord ground truth datasets (p < 0.00002 for all comparisons; Wilcoxon signed-rank tests across neurons within a dataset; see the absolute values in Fig. 5a,c,f,h and Fig. 5-1). However, absolute spike rates were better recovered by retrained CASCADE compared to default CASCADE (p < 0.00001 for both the glutamatergic and the GABAergic datasets; Wilcoxon signed-rank tests; Fig. 5c,d,h,i). Hence, spike rates during high spike-rate events were systematically underestimated by spike rate inference, but less so with a retrained supervised algorithm.

### Spike rate inference of absolute and relative spike rates

Next, we wanted to quantify how accurately overall spike rates across all event types were reflected by spike rate inference with default or retrained CASCADE. First, we compared absolute spike rates estimated by CASCADE with ground truth spike rates and derived the amount of false positives and false negative detection of spiking activity (Fig. 6a). False positive and negative detections depend on the time window of evaluation because a spike event that is correctly detected but shifted in time will be seen as a false detection if a short time window is used for evaluation but as a correct detection if a longer evaluation time window is applied. Accordingly, false detections decreased when the time window of evaluation was increased and converged towards an average bias for the longest time windows (Fig. 6b,c). We found that default CASCADE resulted in a high fraction of false positive detections for the GABAergic dataset and a high fraction of false negative detections for the glutamatergic dataset. Retraining with specific ground truth (again using the “leave-one-out” strategy) improved upon these problems, as is evident from the quantification (Fig. 6d,c). Therefore, retraining not only improved performance across frame rates and noise levels (Fig. 4) but also reduced the bias of absolute inferred spike rates (Fig. 6a-c).

**Figure 6.**
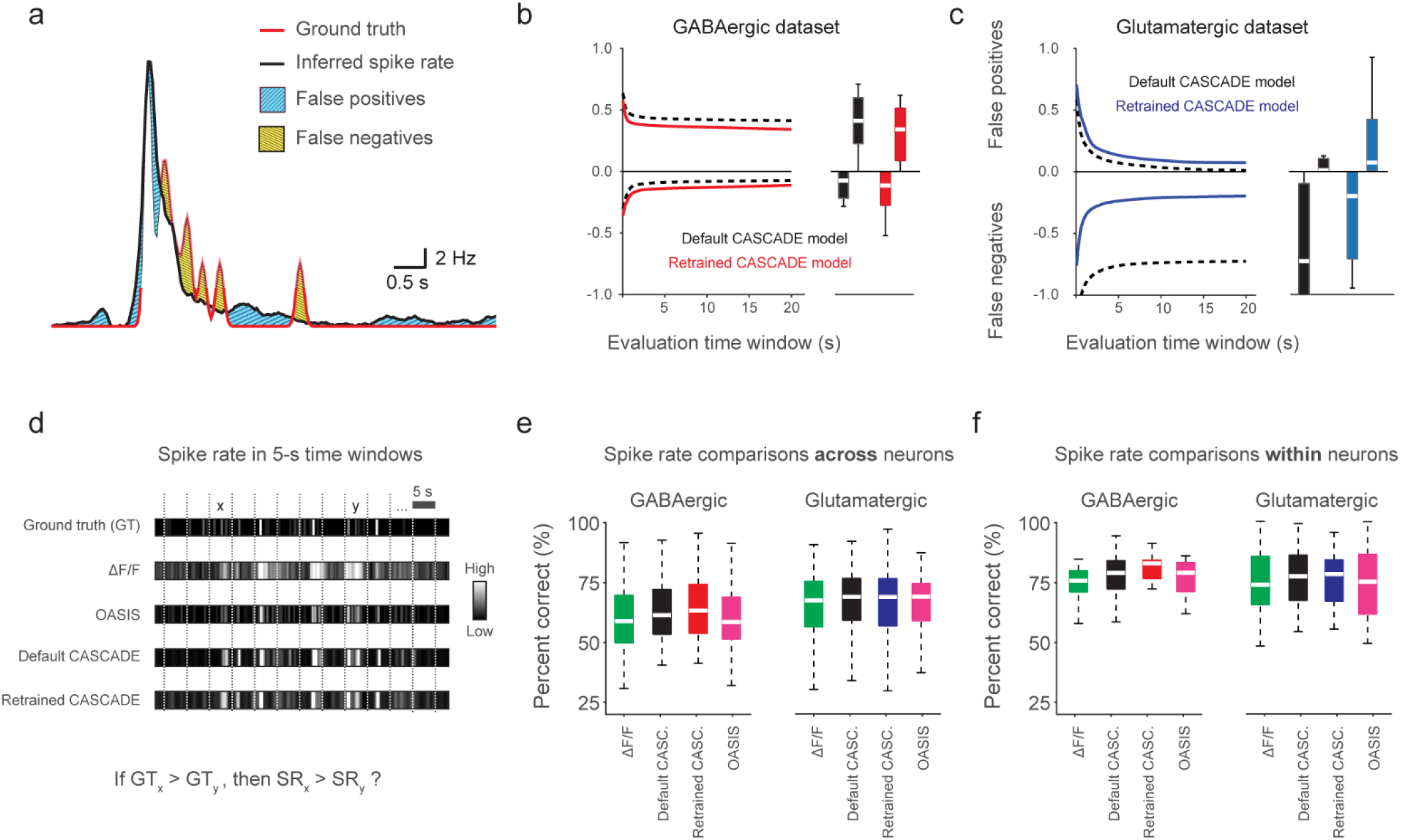
Inference of absolute and relative spike rates with CASCADE. **a**, Illustration of both types of errors, false positives (blue) and false negatives (orange). Ground truth in red, inferred spike rate in black. **b**, False positives and negatives for the default (black) and the retrained (red) SC GABA^+^ CASCADE model. The evaluation time window indicates the binning of inferred spike rates before false positives/negatives were computed as shown in (a). Therefore, the values for the longest evaluation time window indicate the overall bias towards false positives or false negatives. This case is additionally illustrated as a box plot on the right side to provide statistics across neurons (n = 23 neurons). **c**, As in (b), but for glutamatergic neurons (n = 21 neurons) and with the retrained SC GABA^+^ CASCADE model. **d**, Illustration of division of each recording in 5-s segments for ground truth, ΔF/F and inferred spike rate traces. The goal is to test whether inferred spike rate (SR) correctly recovers whether the ground truth spike rate (GT) within a random 5-s segment x is larger than the true spike rate in another segment y. **e**, Percentage of correct spike rate comparisons as described in (d) for both datasets and all approaches. Compared segments are taken from different neurons. Statistics across n = 506 and n = 420 neuron pairs for the GABAergic and glutamatergic datasets. Outcomes of relevant statistical tests are reported in the main text. **f**, Percentage of correct spike rate comparisons as described in (d) for both datasets and all approaches. Compared segments are taken from the same neuron. Statistics across n = 23 and n = 21 neurons for the GABAergic and glutamatergic datasets.

Next, we analyzed how spike rate inference with CASCADE or OASIS improves the estimation of relative spike rates, i.e., the comparison of spike rates, be it across different neurons or for the same neuron across different time windows. To perform this evaluation on the ground truth dataset, we split the recordings into 5-s segments. For each pair of such segments, we evaluated whether the different approaches (raw ΔF/F, OASIS, default CASCADE, retrained CASCADE) correctly identify the segment with the larger number of spikes (Fig. 6d). We performed these analyses first for comparisons of spike rates across neurons, and then for comparisons across 5-s segments within the same neuron.

For segment pairs from different neurons (Fig. 6e), the median percentage of correct evaluations was relatively modest for raw ΔF/F (58% and 67% for the GABAergic and glutamatergic datasets; chance level, 50%), increased modestly after spike rate inference with OASIS (59% and 68%) and default CASCADE (61% and 68%) and further increased for the GABAergic dataset after retraining of CASCADE (63% and 68%; significantly higher than all other approaches for GABAergic neurons, p < 0.001; p < 0.05 compared with ΔF/F and p > 0.05 compared with OASIS and default CASCADE for glutamatergic neurons; one-sided Wilcoxon signed-rank test). For all approaches, a higher spike rate differences between the two compared segments resulted in a higher percentage of correct evaluations, and this correlation was highest for retrained CASCADE (r = 0.33 ± 0.01, mean ± s.d., across 420 neuron-neuron pairs for glutamatergic and 0.39 ± 0.01 across 506 pairs for GABAergic neurons; compared to 0.25 ± 0.01 and 0.30 ± 0.01 for ΔF/F, 0.27 ± 0.01 and 0.33 ± 0.01 for OASIS, 0.32 ± 0.01 and 0.34 ± 0.01 for default CASCADE; all p-values <0.01 for comparisons with retrained CASCADE, one-sided Wilcoxon signed-rank test).

Furthermore, we compared segment pairs within the same neuron (Fig. 6f) and observed an overall much higher percentage of correct evaluations already for ΔF/F (76% and 74% for the GABAergic and glutamatergic datasets), which was further increased after spike rate inference (OASIS: 79% and 75%, default CASCADE: 79% and 77%, retrained CASCADE: 83% and 78%; the latter was significantly higher than all other approaches for GABAergic neurons, p < 0.0005; p > 0.05 for glutamatergic neurons for all comparisons). Again, the correct percentage correlated positively with the spike rate difference between the two segments, and again this correlation was maximal for retrained CASCADE (r = 0.25 ± 0.05, mean ± s.d., across 21 glutamatergic and 0.33 ± 0.03 for 23 GABAergic neurons; compared to 0.18 ± 0.05 and 0.26 ± 0.03 for ΔF/F, 0.18 ± 0.05 and 0.28 ± 0.03 for OASIS, 0.22 ± 0.04 and 0.29 ± 0.03 for default CASCADE; all p-values <0.05 for comparisons with retrained CASCADE). Therefore, taken together, comparisons of spike rates *between* different neurons remain challenging (Fig. 6e), but inference algorithms reliably detect spike rate changes *within* a neuron (Fig. 6f). Nonetheless, spike rate inference, in particular with retrained models, enhanced the statistical power of both comparisons.

### Spike rate inference from anesthetized in vivo spinal cord recordings

Finally, we wanted to test how the CASCADE model retrained on *ex vivo* spinal cord data transferred to *in vivo* imaging conditions. To this end, we analyzed previously acquired *in vivo* spinal cord calcium imaging data from both glutamatergic and GABAergic neurons (Sullivan and Sdrulla, 2022) (Fig. 7a-c). From the extracted ΔF/F traces of the neuronal populations, we performed spike rate inference with dedicated models retrained for the respective neuronal cell types (Fig. 7d,e). Spike rate inference resulted in denoised recordings, most clearly apparent from the cleaned-up baseline for inferred spiking activity from neurons in phases without activity, as expected from previous applications of CASCADE (Rupprecht et al., 2021). In addition, spike rate inference provided an estimate of the number of spikes during a given event (Fig. 7f-h). The most prominent calcium events of a typical duration of 5 s were estimated to be more than 50 spikes (Fig. 7f,g) or even more than 100 spikes, equivalent to a mean spike rate of 10-20 Hz (Fig. 7h). These numbers are consistent with our previous observation that single spikes often evoke indetectable calcium transients; therefore, detectable events are typically associated with a much greater number of spikes (Fig. 3).

**Figure 7.**
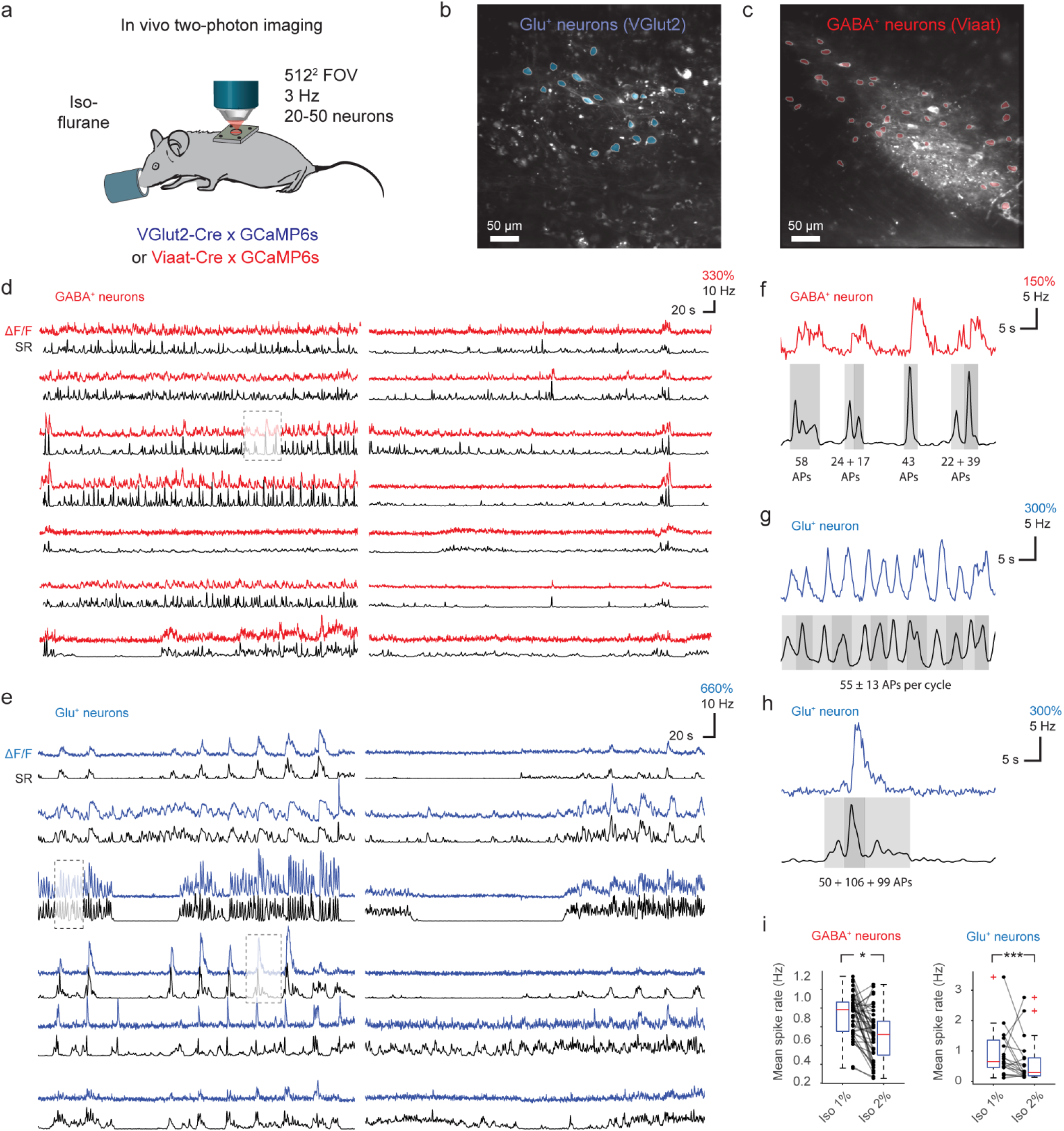
Spike rate inference from population calcium imaging in mouse spinal cord. **a**, Experimental preparation for cell-type specific *in vivo* two-photon imaging of mouse spinal cord in transgenic mice during anesthesia. **b**, Example FOV during imaging of glutamatergic spinal cord neurons. **c**, Example FOV during imaging of GABAergic spinal cord neurons. **d**, Extracted ΔF/F traces (red) and deconvolved spike rates (black) from a subset of the GABAergic neuron ROIs from (c). Deconvolution performed with the SC GABA^+^ CASCADE model. **e**, Extracted ΔF/F traces (blue) and deconvolved spike rates (black) from a subset of the glutamatergic neuron ROIs from (b). Deconvolution performed with the SC Glu^+^ CASCADE model. **f**, Zoom-in to a ΔF/F trace and estimated spike rate for a GABAergic neuron in (d). The number of spikes (action potentials, “APs”) is indicated below events. Subevents are shaded with different gray values. **g-h**, Same as (f) but for glutamatergic neurons in (e). **i**, Mean spike rates of neurons detected from the FOVs in (b,c) during shallow (1% isoflurane) vs. deep (2%) anesthesia, with decreased spike rates for deep anesthesia for both cell types (p = 0.040, Wilcoxon ranksum test, 23 neurons for the glutamatergic dataset; p = 0.000016, 48 neurons for the GABAergic d1a7taset). *p < 0.05, **p < 0.01, ***p < 0.001, n.s., not significant (p > 0.05).

Furthermore, to test if retrained CASADE could detect *in vivo* changes in activity within individual neurons, the level of isoflurane anesthesia was adjusted to either 1% or 2%. As expected, an increase of anesthesia depth substantially decreased the inferred spike rates for the majority of neurons (Fig. 7i). This application showcases that the retrained CASCADE models can be used to estimate spike rates and denoise *in vivo* recordings.

## Discussion

In this study, we investigated how information about spike rates can be recovered from calcium imaging data in mouse spinal cord. We recorded ground truth datasets with simultaneous calcium imaging and electrophysiology in genetically identified glutamatergic and GABAergic spinal cord neurons. Based on these datasets, we trained a supervised deep learning algorithm that infers spike rates from calcium imaging data, and we provide these pretrained models in an easy-to-use framework (CASCADE). The original CASCADE model, trained with data from cortical neurons, generalized well to these spinal cord neurons, but specific retraining on spinal cord ground truth improved the retrieval of high-frequency spike events, was less biased in predicting absolute spike rates and resulted in an improved prediction of relative spike rates across and within neurons.

Spike inference algorithms can be evaluated based on two qualitatively different outputs, either discrete inferred spike times (Deneux et al., 2016; Hoang et al., 2020; Lütcke et al., 2013; Oñativia et al., 2013; Pnevmatikakis et al., 2016) or continuous and smooth spike rates (Berens et al., 2018; Rupprecht et al., 2021; Theis et al., 2016; Zhou et al., 2023). Here, we only used the latter approach since single spikes cannot be identified under our typical recording conditions (Fig. 3a-d). We used the correlation with true spike rates as our main metric and found that supervised spike rate inference with CASCADE yielded substantially better results than spike rate inference with OASIS or by using raw ΔF/F as a proxy for spiking activity (Fig. 4). This performance gap was reduced when we decreased the temporal precision used to evaluate the algorithms (Fig. 4g; Fig. 4-3c). These analyses indicate that raw ΔF/F or spike rate inference with OASIS are a good proxy for spiking activity when temporal precision is neglected, while supervised spike rate inference with CASCADE performs well for both low and high temporal precision. Overall, a default version of CASCADE, trained on a large database across diverse brain regions but not including the spinal cord, performed very well on the before unseen spinal cord data. This result highlights the robustness of spike rate inference across CNS regions, and supports previous analyses demonstrating generalization of spike rate inference across datasets (Rupprecht et al., 2021).

However, we also observed limitations of this capability to generalize to spinal cord data. To improve upon the default algorithm, we performed ground truth recordings in glutamatergic and GABAergic spinal cord neurons. It has been previously shown that a much larger or smaller spike-evoked calcium transient can make it challenging to generalize across datasets, e.g., from excitatory neurons to fast-spiking interneurons (Ali and Kwan, 2020; Rupprecht et al., 2021). Our ground truth recordings show that spike-evoked calcium transients are, on average, similar between the GABAergic and glutamatergic spinal cord populations (Fig. 3a-c), with similar decay kinetics. This is noteworthy since several interneurons in the brain have been shown to exhibit much lower spike-evoked calcium transients than cortical pyramidal cells (Aponte et al., 2008; Kwan and Dan, 2012).

In addition, we observe that spike-evoked calcium transients from our spinal cord dataset displayed slower decay times but similar amplitudes compared to recordings from mouse cortex (Fig. 3a). The comparable amplitude lends further support to the transferability of spike rate inference from cortical to spinal cord recordings. It must, however, be kept in mind that our *ex vivo* ground truth recordings are only an approximation of *in vivo* conditions, as the spinal cord is resected, perfused with room temperature ACSF and deafferented. Although we did not observe an effect of room vs. physiological temperature recordings on the spike-evoked calcium transient (Fig. 3-1), we cannot exclude potential temperature-related effects with certainty. Furthermore, using the GCaMP signal to target cells for our cell-attached recordings may have introduced a bias toward sampling brightly visible cells, which tend to exhibit higher spontaneous activity. *In vivo* ground truth recordings in spinal cord are challenging due to the difficulty of access and due to prominent motion artifacts (Johannssen and Helmchen, 2010; Nelson et al., 2019; Sullivan and Sdrulla, 2022). Hence, our ground truth recordings represent a compromise that matches the natural conditions as much as possible. In support of this interpretation, calcium signals of ground truth recordings (Fig. 1) were of similar ΔF/F amplitude compared to previous in vivo recordings (Sullivan and Sdrulla, 2022), and their spike rates (Fig. 2) were of the same order of magnitude as for electrophysiological recordings (Lucas-Romero et al., 2018) and as the spike rates inferred from anesthetized in vivo recordings (Fig. 7). In addition, spike patterns in our ground truth recordings included both bursts and more regular spiking patterns, in agreement with observations *in vitro* and *in vivo* across multiple species both for spontaneous and stimulus-evoked patterns (Kumazawa and Perl, 1978; Lucas-Romero et al., 2022, 2018; Medrano et al., 2016; Sandkühler and Eblen-Zajjur, 1994). A caveat to keep in mind are the diverging definitions of specific spike patterns across fields. For example, in this study, we investigated electrophysiologically defined bursts (inter-spike interval <10 ms; Fig. 2), but also high-frequency spike events (peak spike rate of >45 Hz; Fig. 5). These event types overlap but do not coincide with the definition of “bursty” spike patterns defined in previous studies on the spinal cord (Lucas-Romero et al., 2022, 2018). Irrespective of these definitions, our dataset (>70,000 spikes over 7.4 hours) includes a diversity of firing patterns and is therefore the best available dataset to train a supervised spike inference algorithm for spinal cord neurons.

In general, retraining a supervised spike inference algorithm with a specific ground truth dataset can enable more precise interpretations of population imaging data. Here, we found that spike rate inference with a retrained version of CASCADE was improved compared to OASIS but also compared to default CASCADE (Fig. 4). Interestingly, this improvement as measured by correlation with ground truth was more pronounced for GABAergic compared to glutamatergic spinal cord neurons (Fig. 4d,e), probably reflecting the higher similarity between glutamatergic spinal cord neurons with (glutamatergic) pyramidal cells from cortex. In addition, we found that high-frequency spiking events were temporally broadened by OASIS but also by default CASCADE (Fig. 5), presumably because both approaches had been optimized for cortical datasets (Fig. 2) and for different calcium kernel decay times (Fig. 3a). For retrained CASCADE, on the other hand, the temporal confinement of spikes during high-frequency spike events as well as their absolute number was more accurately recovered. This absolute number of spikes during such events was still underestimated, as found similarly before (Rupprecht et al., 2021), most likely due to saturation of the calcium indicator. Furthermore, retraining of CASCADE resulted in a less biased estimate of absolute spike rates for both GABAergic and glutamatergic neurons (Fig. 6a-c). Nonetheless, all spike rate inference models proved useful in detecting relative changes in spinal cord neuron spiking, but retrained CASCADE performed slightly better. This performance improvement most likely reflects the previously described suppression of noise by spike rate inference (Pachitariu et al., 2018; Rupprecht et al., 2021). Together, these results highlight the additional benefits in terms of temporal precision and absolute calibration by retraining a supervised algorithm on specific ground truth data. We provide pretrained spike rate inference models for glutamatergic and GABAergic spinal cord neurons together with the underlying ground truth in an easy-to-use repository (https://github.com/HelmchenLabSoftware/Cascade).

An important observation of our analysis is that spike rates can be readily compared for different temporal segments of the same neuron but not equally well across neurons (Fig. 6d-f). This is not an artifact introduced by spike rate inference, since the effect was similarly present in ΔF/F data. A potential explanation of this problem is the large variability of spike-triggered ΔF/F transients across neurons within the same dataset, both for spinal cord and cortex (Fig. 3b,c). This suggests that comparisons of spike rates across neurons may be unreliable, not only for spinal cord but also cortical data. The origin of this variability is unclear, but it may be due to variable levels of indicator concentrations across cells (Éltes et al., 2019). An alternative and not mutually exclusive explanation is that the determination of the baseline level F_0_ to compute ΔF/F_0_ is error-prone due to the low baseline fluorescence of GCaMP, thereby inducing biases dependent on bleed-through from surrounding neuropil. Independent of its origin, this variability constitutes a limitation for the inference of absolute spike rates and for the comparison of spiking activity across neurons. This problem has been addressed before with attempts to autocalibrate ΔF/F in order to identify the unitary response amplitude without ground truth (Deneux et al., 2016; Éltes et al., 2019). While these approaches have so far not been commonly used due to the ambiguity of autocalibration, our study emphasizes the need for improving on such tools to enable quantitative inference of absolute spike rates. We hope that our results and the openly shared CASCADE inference models will encourage future work to calibrate calcium data in other brain regions and to address the remaining limitations due to the variability of spike-triggered calcium responses across neurons.

## Contributions

P.R. and A.S. conceived the study. W.F., S.S. and A.S. performed all experiments and pre-analyses. P.R. and A.S. performed quality control. P.R. wrote code, performed analyses and visualized results. P.R., S.S., F.H. and A.S. wrote the paper. F.H. and A.S. supervised the project.

## Acknowledgements

This work was supported by the National Institutes of Health (Grants K08NS099503, RF1NS124557 to A.S.) and by grants from the Swiss National Science Foundation (project grant 310030B_170269 to F.H.; Ambizione grant PZ00P3_209114 to P.R.).

## Extended Data Figures

**Figure 1-1.**
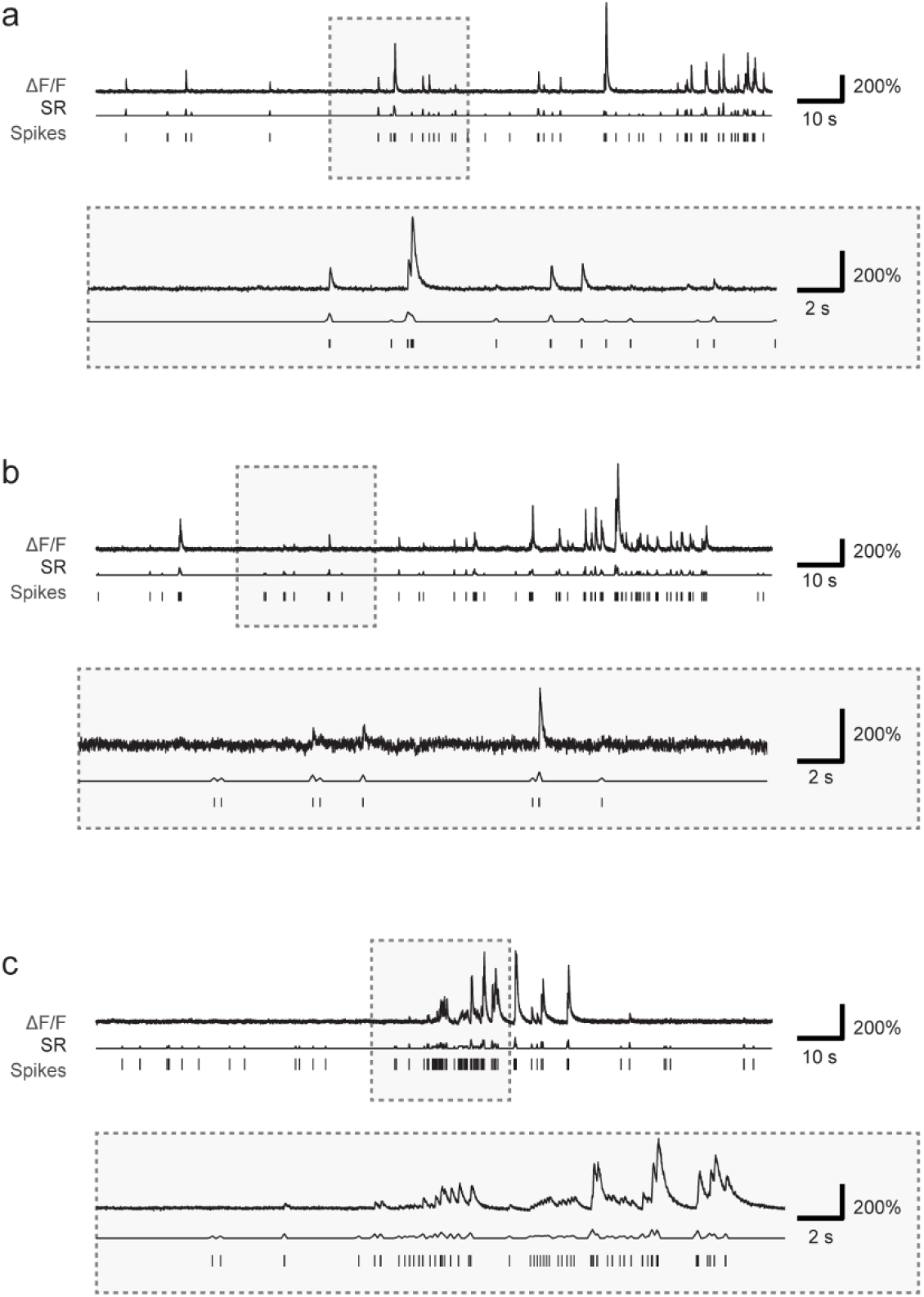
Figure Simultaneous recording of action potentials and calcium signals in layer 2/3 pyramidal neurons from mouse cortex. Same presentation of example recordings as in Fig. 1e-j but for the ‘cortex dataset’, comprising ground truth recordings from four different transgenic mouse lines expressing GCaMP6f or GCaMP6s in layer 2/3 neurons in mouse visual cortex (Huang et al., 2021). *ΔF/F*, normalized fluorescence extracted from calcium imaging; *SR*, smoothed spike rate derived from electrophysiological spike times; *Spikes*, spike times detected from the electrophysiological recording.

**Figure 3-1.**
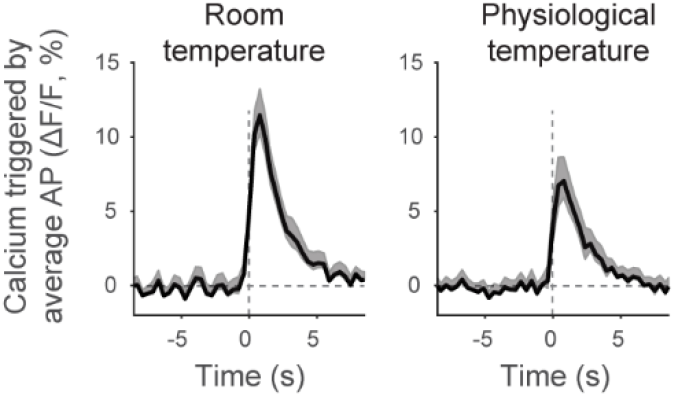
Temperature-dependence of the calcium transient evoked by the average spike. Calcium response (ΔF/F) for the average action potential across neurons, computed by linear deconvolution of the ground truth recording. *Left*: All neurons from the *SC Glu*^*+*^ dataset. *Right*: Recordings from the *SC Glu*^*+*^ dataset (11 out of 69 recordings in 5 out of 21 neurons) that were performed at physiological temperature (37°C). No slowing of indicator kinetics for room temperature is visible from this dataset (single exponential fit: τ = 2.9 ± 0.5 s for room temperature, 3.0 ± 0.6 s for physiological temperature; fit ± 90% confidence intervals).

**Figure 4-1.**
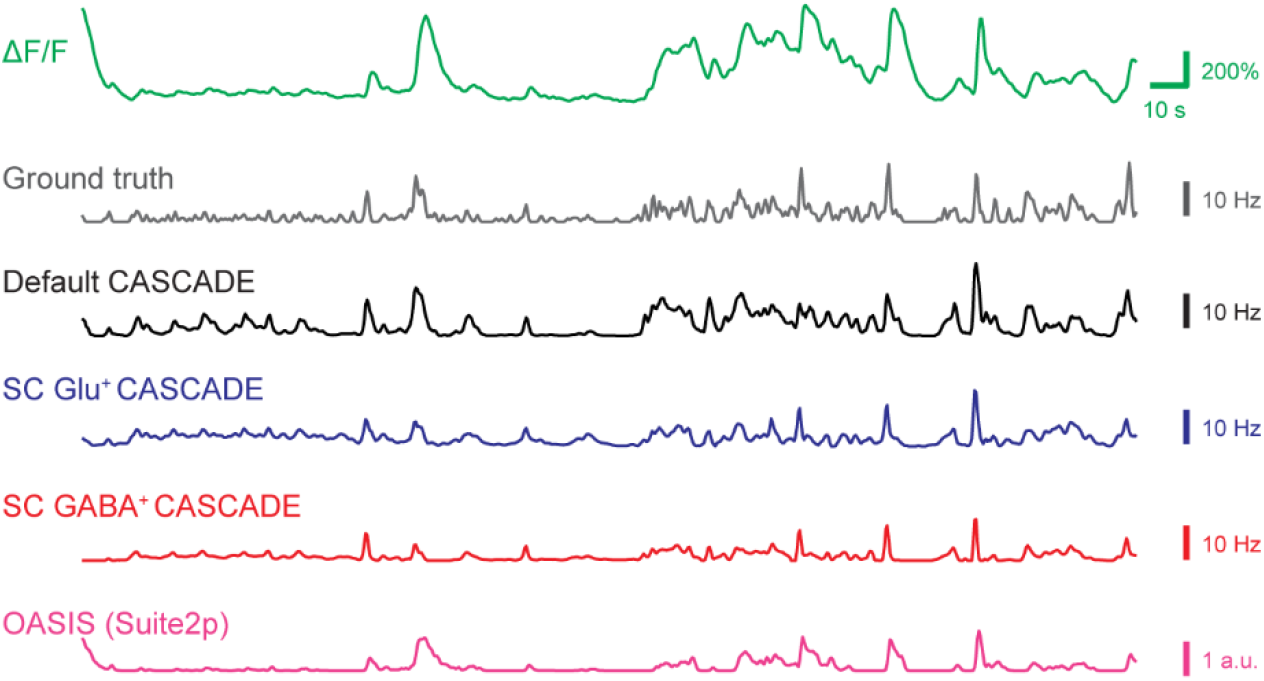
Example predictions for a lower sampling rate. Examples of an extracted ground truth recording (same as in Fig. 4a) together with the ground truth spike rate and spike rate predictions with CASCADE and OASIS. Sampled at 2.5 Hz with a standardized noise level of “7”.

**Figure 4-2.**
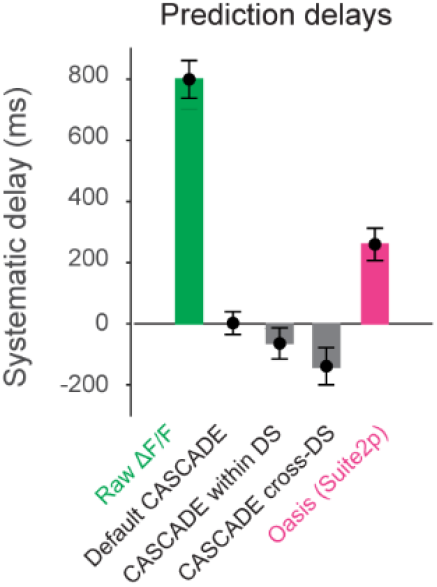
Systematic delay of spike rate inference with respect to ground truth. Quantified for the approaches described in Fig. 4a. “Within-DS” indicates that CASCADE was trained with matching datasets (*e*.*g*., CASCADE trained on glutamatergic neurons and applied to glutamatergic neurons), while “cross-DS” was trained with non-matching datasets. Consistent with previous analyses (Rupprecht et al., 2021), not only raw ΔF/F but also OASIS tended to result in a systematic delay of predictions compared to ground truth, as opposed to default or retrained CASCADE. All analyses in Fig. 4 to 7 are corrected for these systematic delays.

**Figure 4-3.**
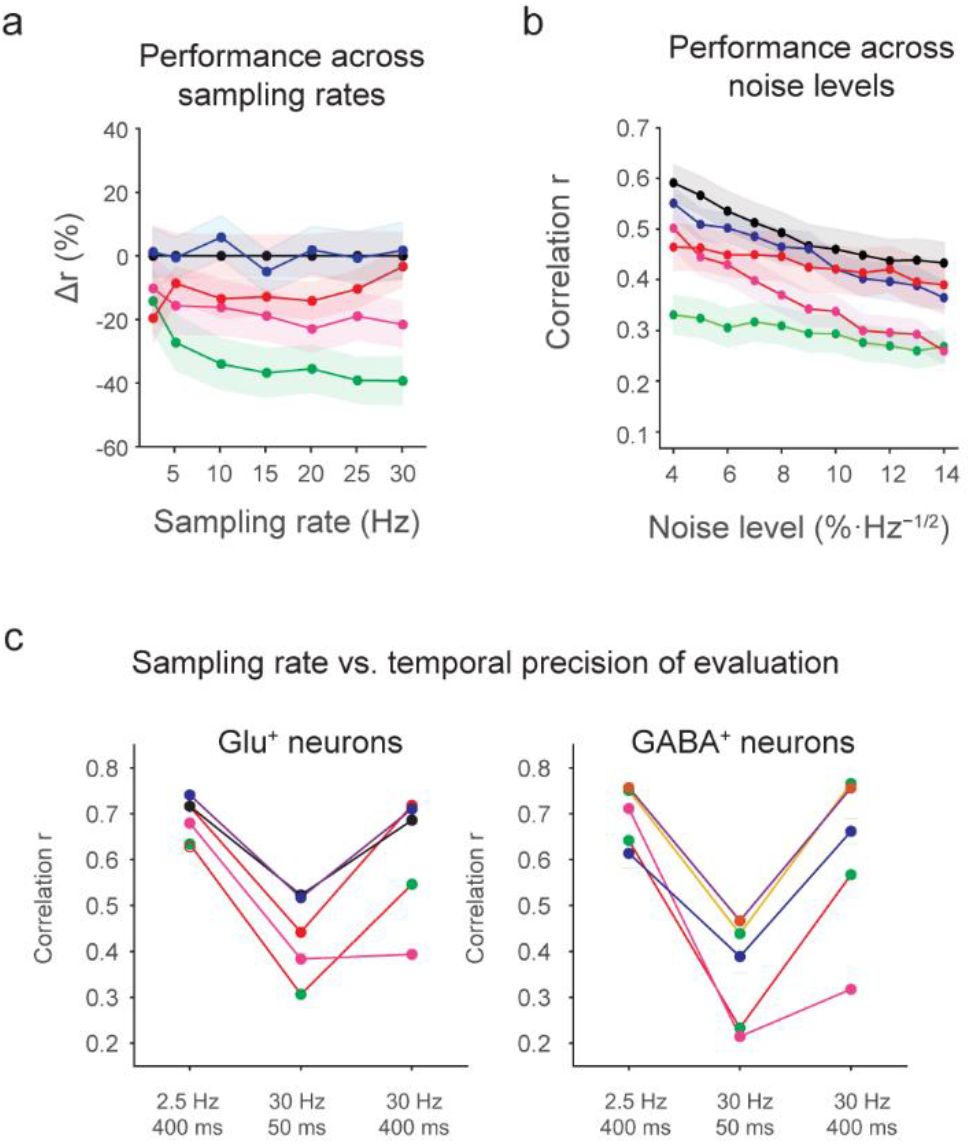
Further comparison of the performance of spike rate inference across algorithms. **a**, Quantification of performance for the glutamatergic spinal cord dataset across re-sampled imaging rates. The same quantification for the GABAergic spinal cord dataset was shown in Fig. 4g. **b**, Quantification of performance for the glutamatergic spinal cord dataset across noise levels. The same quantification for the GABAergic spinal cord dataset was shown in Fig. 4h. **c**, Control analysis to show that performance (“correlation”) is recovered for high imaging rates when the evaluation criterion (temporal smoothing applied before correlation with ground truth) is matched to low imaging rates. For example, the performance underlying the left-most datapoints were measured at an imaging rate of 2.5 Hz with a smoothing window of 400 ms. Therefore, spike rate inference reconstructs spike rates with similar accuracy from fast and slow imaging data when evaluated with the same slow temporal precision.

**Figure 4-4.**
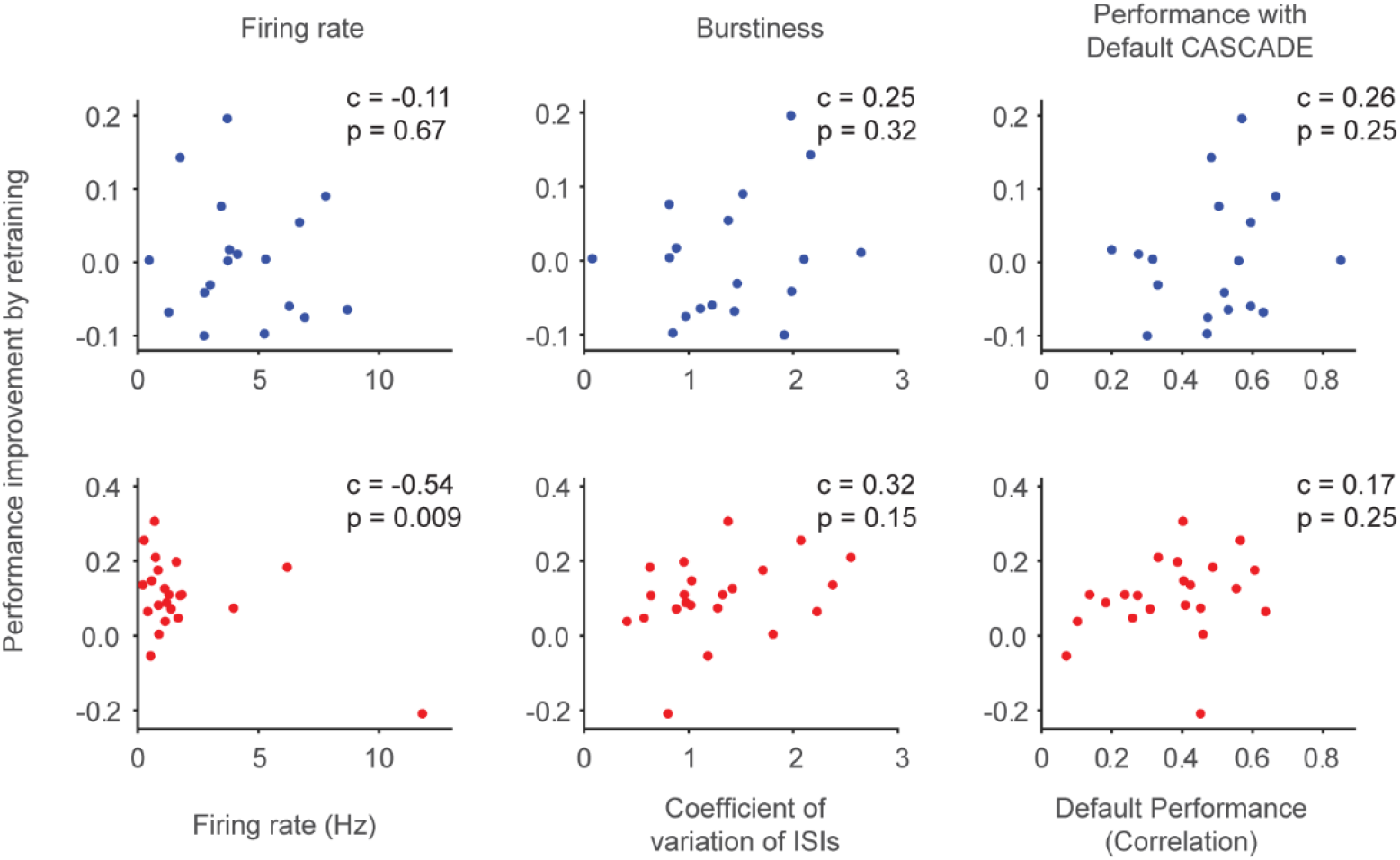
Dependence of performance improvement on cellular characteristics. To test whether specific subtypes of spinal cord dorsal horn neurons improved more than others with retraining, we explained the improvement by retraining (y-axis) by other variables obtained for each cell (average firing rate, burstiness, see Fig. 2; and the performance when applying Default CASCADE). Top row: glutamatergic neurons. Bottom row: GABAergic neurons. We did not find any significant correlation (p > 0.05; correlation values c and significance values p indicated in the figure) for either excitatory (blue) or inhibitory (red) spinal cord neurons. The relationship between firing rate and performance improvement was statistically significant for inhibitory neurons (p = 0.009), but this effect was driven by a single outlier (p = 0.90 after removal of the single outlier). As a conclusion, no cellular properties potentially indicative of cellular subtypes in the spinal cord were found that were predictive of performance improvement after retraining. It is therefore reasonable to assume that, within the limitations of these limited datasets, performance improvements upon retraining of CASCADE affected most or all neurons without a specific pattern.

**Figure 5-1.**
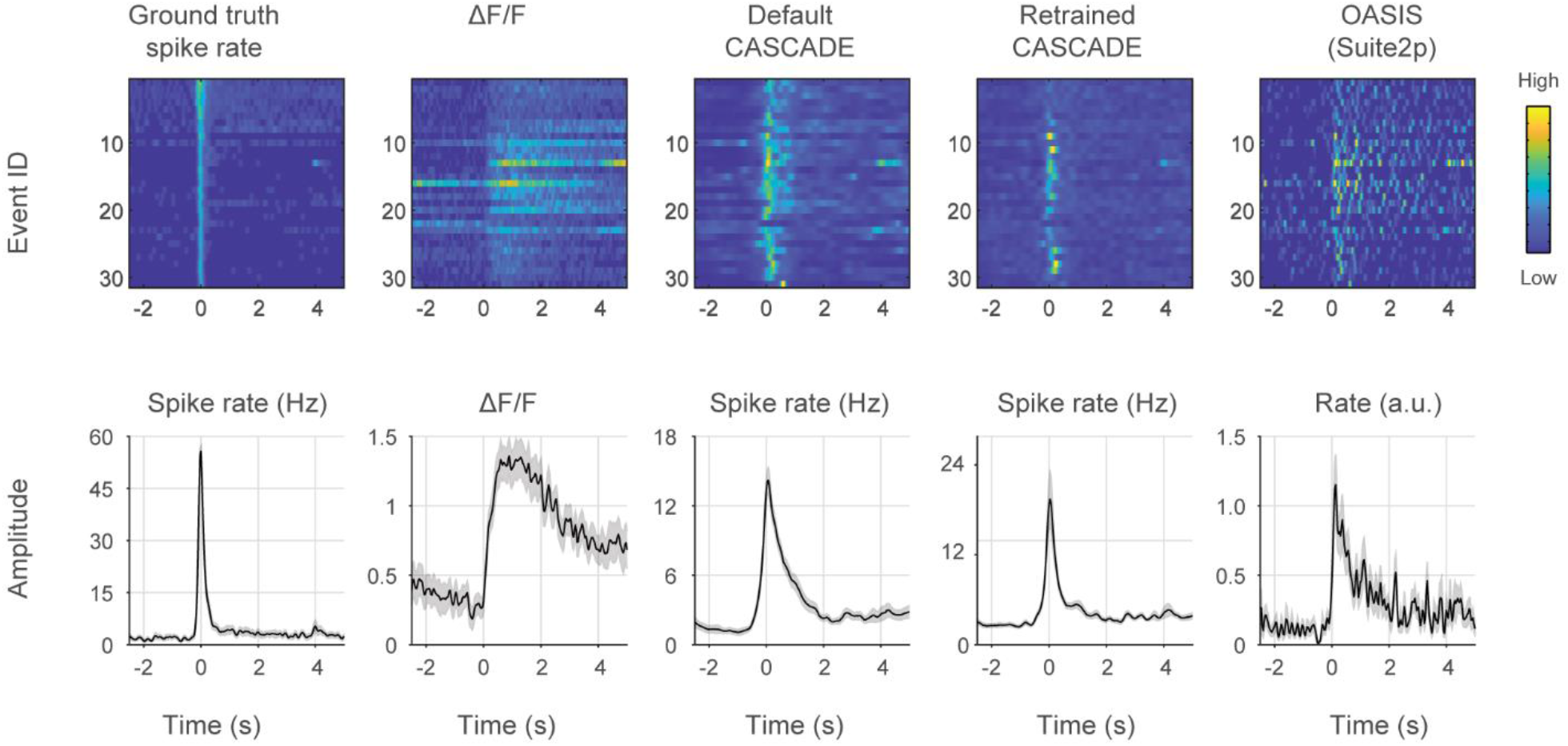
Spike rate inference for high-frequency spike events for GABAergic neurons in spinal cord. *Top row:* High-frequency spike events for the GABAergic neuron dataset, with the corresponding associated ground truth spike rate, ΔF/F signal, spike rate inferred by the default CASCADE model, spike rate inferred from the retrained CASCADE model, and spike rate inferred by the OASIS algorithm. *Bottom row:* Same as in top row panels but averaged across events, with absolute values indicated if possible. Prolonged spike rate is seen for the default CASCADE and OASIS models. The median number of spikes during events (1-s window around event) for ground truth vs. default CASCADE vs. retrained CASCADE is 17.2 vs. 10.0 vs. 11.8 spikes.

## Notes

https://github.com/HelmchenLabSoftware/Cascade

### Competing Interest Statement

The authors have declared no competing interest.

### Summary of Updates

Error in Figure 3 (wrong scaling) fixed; additional control analyses performed (Fig. 4-4); improved wording of Discussion and Methods; title adapted ("spike rate inference" instead of "spike inference")

